# Crop management shapes the diversity and activity of DNA and RNA viruses in the rhizosphere

**DOI:** 10.1101/2022.04.22.488307

**Authors:** George Muscatt, Sally Hilton, Sebastien Raguideau, Graham Teakle, Ian D. E. A. Lidbury, Elizabeth M. H. Wellington, Christopher Quince, Andrew Millard, Gary D. Bending, Eleanor Jameson

## Abstract

**Background:** The rhizosphere is a hotspot for microbial activity and contributes to ecosystem services including plant health and biogeochemical cycling. The activity of microbial viruses, and their influence on plant-microbe interactions in the rhizosphere, remains undetermined. Given the impact of viruses on the ecology and evolution of their host communities, determining how soil viruses influence microbiome dynamics is crucial to build a holistic understanding of rhizosphere functions.

**Results:** Here, we aimed to investigate the influence of crop management on the composition and activity of bulk soil, rhizosphere soil, and root viral communities. We combined viromics, metagenomics, and metatranscriptomics on soil samples collected from a 3-year crop rotation field trial of oilseed rape (*Brassica napus* L.). By recovering 1,059 dsDNA viral populations and 16,541 ssRNA bacteriophage populations, we expanded the number of underexplored *Leviviricetes* genomes by > 5 times. Through detection of viral activity in metatranscriptomes, we uncovered evidence of “Kill-the-Winner” dynamics, implicating soil bacteriophages in driving bacterial community succession. Moreover, we found the activity of viruses increased with proximity to crop roots and identified that soil viruses may influence plant-microbe interactions through the reprogramming of bacterial host metabolism. We have provided the first evidence of crop rotation-driven impacts on soil microbial communities extending to viruses. To this aim, we present the novel principal of “viral priming”, which describes how the consecutive growth of the same crop species primes viral activity in the rhizosphere through local adaptation.

**Conclusions:** Overall, we reveal unprecedented spatial and temporal diversity in viral community composition and activity across root, rhizosphere soil and bulk soil compartments. Our work demonstrates that the roles of soil viruses need greater consideration to exploit the rhizosphere microbiome for food security, food safety, and environmental sustainability.

## Background

Soils harbour organisms from multiple kingdoms of life and provide ecosystems for > 25% of Earth’s biodiversity [1]. Viruses, the smallest microorganisms in terrestrial ecosystems, often exceed the number of co-existing bacteria [2], with up to 10^10^ virus-like particles per gram of soil [3]. Of particular interest are the viruses of microbes, whose lytic activity provides top-down control of microbial host populations, and whose expression of viral encoded auxiliary metabolic genes (AMGs) modulates host metabolism [4–7]. In marine ecosystems viruses have been estimated to turnover ∼ 20% of microbial biomass each day [8], resulting in drastic impacts on ocean carbon and nutrient cycling [9, 10]. Given that there is an estimated 70 times more terrestrial biomass than marine biomass [11], and that viral infection rates are speculated to be greater in soils than oceans [12], there is significant interest in unearthing the importance of viruses in terrestrial ecosystems [13, 14]. The physical structure of soil, however, hinders the extraction and subsequent cultivation of soil viruses, resulting in the current knowledge gap surrounding the ecological roles of viruses in soils [15].

Circumventing the requirement to culture viruses and their microbial hosts, metagenomics and viral size-fractionated metagenomics (viromics), have facilitated the estimation of total viral community composition and diversity across Earth’s ecosystems [16–20]. Moreover, the recent optimisation of soil viromics protocols [15, 21, 22] and *de novo* viral prediction tools [23–25] have enabled the systematic characterisation of soil viral communities. However, conventional DNA approaches are unable to reveal the activity of recovered viruses. To overcome this, metatranscriptomics can be applied to characterise gene expression through the quantification of sequenced messenger RNA transcripts [26]. Given that viruses require host cell machinery for the transcription of their genes, viral activity can be used to indicate host infection. Additionally, RNA viral genomes can be assembled from metatranscriptomes, which has revealed the abundance and activity of single-stranded RNA (ssRNA) bacteriophages (phages) in both non-terrestrial [27–29] and terrestrial ecosystems alike [30–33]. For example, the discovery and emergent role of a disproportionately understudied class of ssRNA soil phages in terrestrial biogeochemistry, named *Leviviricetes* [30, 31]. Despite the advantages of combining metatranscriptomics with metagenomics to simultaneously investigate the composition and activity of DNA and RNA viral communities, there has been no such implementation in previous soil viromics studies.

Plants release ∼20% of the carbon assimilated during photosynthesis into the soil through root exudates [34]. This provides labile nutrients and energy to microorganisms in the soil adjacent to the root system, known as the rhizosphere. Subsequently, the rhizosphere soil compartment contains greater microbial density and activity than surrounding bulk soil [35], and contributes to ecosystem services including plant health and biogeochemical cycling [36–38]. While growing evidence implicates soil viruses in contributing to terrestrial carbon and nutrient cycling [16, 39–43], viruses remain a black box in soil and rhizosphere ecology. It is unclear whether the rhizosphere is a zone of high viral density and activity, and the effects of viral activity on plant-microbe interactions remain undetermined.

Agricultural management utilises a variety of strategies to maintain soil fertility and productivity. The impacts of these on soil and rhizosphere microbiomes have been intensively studied, with the focus on prokaryote and eukaryote communities [44, 45], while interactions with viral communities have received little or no attention. Crop rotation is a widespread practice in which different crop plant species are grown sequentially to improve soil fertility and reduce pest and pathogen pressures [46, 47]. Subsequently, crop rotation has been associated with shifts in bacterial, fungal, and archaeal community compositions, with resulting benefits to crop health and yield [48–51]. However, there is no understanding of the effects of rotation on viral communities, or the associated interactions with microbial communities. Given the impact of viruses on the ecology and evolution of their host communities in non-soil systems [52, 53], determining the roles of soil viruses in moderating microbiome dynamics is crucial to build a holistic understanding of rhizosphere functions [54].

Thus, we aimed to investigate the influence of crop rotation on the composition and activity of bulk soil, rhizosphere soil, and root viral communities. Combining viromics, metagenomics, and metatranscriptomics, we recovered novel double-stranded DNA (dsDNA) and ssRNA viral operational taxonomic units (vOTUs), expanding the number of known *Leviviricetes* genomes by > 5 times. Next, we simultaneously estimated the relative contributions of compartment and crop rotation in shaping the composition of DNA and RNA soil viral communities, relative to bacterial communities. Lastly, we characterise the spatiotemporal activity of DNA viral communities across three stages of crop growth, revealing dynamic viral-host interactions across the root-associated microbiomes.

## Methods

### Field site

The field site was established in 2014 at the University of Warwick Crop Centre in Wellesbourne, UK, following conventional management, as previously described [51]. 8 plots of 24 m × 6 m were set up as shown in **Fig. S1**, allowing for 4 replicate samples of the two crop management practices. Two crop growth strategies of oilseed rape (*Brassica napus* L.) were adopted: continuous cropping, whereby oilseed rape was grown for three consecutive years; and virgin rotation, whereby oilseed rape was grown following two preceding years of winter wheat (*Triticum aestivum*). The soil was a sandy-loam of the Wick series, with 73% sand, 12% silt, 14% clay, a pH of 6.5 and organic carbon content of 0.8% [55].

### Sample collection

Samples were collected from each plot at three time points (November 2016, March 2017, June 2017) during the growing season of year 3 (2016/2017). For each sample, eight plants were taken from the plot by sampling ∼1 m into the plot to avoid the edge. Loosely adhered soil was removed from the roots by tapping. The roots from all eight plants (or for large roots, 6 × ∼5 cm root sections were used per plant) were transferred to a 50 mL tube containing 20 mL autoclaved Milli-Q water and shaken for 20 s (first wash). The roots were transferred to a second tube and washing was repeated (second wash). The first and second washes were combined and frozen in liquid nitrogen (rhizosphere soil samples). The roots were washed a final time in 20 mL water, transferred to an empty 50 mL tube, and frozen in liquid nitrogen (root samples). This whole process was performed in the field in < 5 mins. Bulk soil was sampled from each plot by selecting areas between plants ∼ 50 cm into the plot to avoid the edge. 2-3 mm of surface soil was removed, and an auger was used to collect soil to a depth of 15 cm. 8 soil cores were sampled per plot, combined, and added to a falcon tube containing 30 mL water to take the total volume to 45 mL. The tubes were shaken for 40 s and frozen in liquid nitrogen (bulk soil samples). All samples were stored at −80°C. The rhizosphere soil and bulk soil samples were subsequently freeze-dried. Root samples were homogenised under liquid nitrogen using a mortar and pestle.

### RNA and DNA extractions

RNA extractions were performed on all root and soil samples from the three time points. RNA was extracted from 1 g of homogenized root or 2 g of soil (rhizosphere soil or bulk soil) using the RNeasy PowerSoil Total RNA Isolation Kit (Qiagen, Hilden, Germany) with two homogenisations in a Fastprep machine (MP Biomedicals) at 5.5 m/s for 30 s, resting on ice for 5 mins between runs. RNA was eluted in 50 µL elution buffer and 46 µL was subsequently DNase treated (DNase Max™) according to the manufacturer’s instructions. The DNase was then removed using the DNase Max™ Removal Resin. The RNA was checked for residual contaminating DNA using 16S rRNA gene universal primers. DNA-free RNA was then purified using RNAClean XP Beads (New England Biolabs) according to the manufacturer’s instructions. The RNA was quantified using Qubit RNA BR kit on a Qubit® fluorometer (Invitrogen, CA, USA).

DNA extractions for total metagenomes and size-fractionated metagenomes (DNA viromes) were only performed on soil samples from the second time point (March 2017). For these samples, total DNA was eluted from the same column as the RNA extractions using the RNeasy PowerSoil DNA Elution Kit (Qiagen, Hilden, Germany), according to the manufacturer’s instructions. The DNA was quantified using Qubit DNA HS and its purity profile checked using a Nanodrop 2.

Size-fractionated DNA for DNA viromes was extracted from ∼ 5 g of soil. Briefly, soil was mixed into a total volume of 50 mL of sterile PBS and shaken vigorously for ∼ 5 mins, before being gently agitated on a tube roller for 1 hour. Following centrifugation at 500 g to pellet large material, the supernatant was removed, sequentially filtered through 0.44 µm and 0.22 µm pore size filters and concentrated using Amicon 100 kDa columns as previously described. The sample was DNAse I treated (1 U/µL) for 60 min at room temperature to remove free contaminating DNA. Viral fraction DNA was extracted through sequential rounds of phenol: chloroform as previously described [56].

### Library construction and sequencing

RNA sequencing was performed by the Earlham Institute, Norwich, UK. Libraries were made using the Illumina TruSeq RNA library (HT, non-directional) kit and all libraries were run across two lanes of the Illumina HiSeq 2500 platform (2 x 150 bp). Following sequencing, Trimmomatic v0.36 [57] was used to remove any TruSeq adapters from the sequences. SortmeRNA [58] was then used to separate and retain the rRNA reads. The forward reads (R1) were quality filtered using VSEARCH with a fast-maxee of 1 and a minimum length of 100 nt. This dataset was used as the raw metatranscriptome for read mapping.

The 16S rRNA gene operational taxonomic unit (OTU) table was generated by first assigning taxonomy to rRNA reads using QIIME and the SILVA database (version 132) at 99% identity. Then, only reads assigned to bacteria (representing 16S rRNA gene transcripts) were retained, while reads assigned to mitochondria or chloroplasts were removed.

Libraries for total metagenome sequencing were prepared and sequenced by Novogene Ltd on an Illumina HiSeq (2 x 150 bp).

Libraries for DNA virome sequencing were prepared using 1 ng of input DNA for the NexteraXT library preparation, following the manufacturer’s instructions. Libraries were sequenced on an Illumina MiSeq in 2 flow cells using v3 chemistry (2 x 300 bp).

### Read processing and assembly

Metatranscriptome reads were quality filtered and trimmed with trim_galore v0.5.0_dev, and then assembled with SPAdes v3.14.0 [59, 60] using the script rna.spades.py and default settings. Total metagenome reads were quality filtered and trimmed with trim_galore v0.5.0_dev [61], and then assembled with MEGAHIT v1.2.9 [62, 63] using –kmer steps of “27,37,47,57,67,77,87,97,107,117,127,137,141”. DNA virome reads were quality filtered and trimmed with sickle v1.33. Viral DNA libraries were then assembled with MEGAHIT using – kmer steps of “21,41,61,81,101,121,141,161,181,201,221,241,249”.

### Recovery of viral populations

dsDNA viral contigs were predicted from the pooled assembled reads from all soil samples, independently for each of the three libraries, i.e., DNA virome, total metagenome and metatranscriptome. For the DNA virome and metatranscriptome, viral contigs were predicted with DeepVirFinder v1.0 [25] and filtered for *q* < 0.05 (estimated for false discovery rate of 0.1) and contig length ≥ 10 kb. For the total metagenome, viral contigs were predicted with VIBRANT v1.0.1 [24] and filtered for contig length ≥ 10 kb, with proviral sequences > 5 kb retained. dsDNA viral contigs predicted from the three libraries were combined and de-duplicated at 95% nucleotide identity across 95% of the contig length using CD-HIT v4.6 [64] to define 1,059 non-redundant vOTUs, representing approximately species-level dsDNA viral populations, in accordance with benchmarking [65]. To determine whether any recovered vOTUs represented previously isolated phage species, we computed the pairwise MinHash genome distances (*D*) to a custom database of all complete phage genomes that were available at the time (May 2020) [66] using MASH v2.0 [67]. Average nucleotide identity (ANI) was estimated by 1 − *D*, and two genomes with ANI values ≥ 95% were considered to represent the same species.

Positive-sense ssRNA phage contigs were predicted from the pooled assembled metatranscriptome reads for each soil sample [27]. The resulting contigs were de-duplicated at 100% global identity using CD-HIT to identify 187,588 non-redundant ssRNA phage contigs, representing 16,541 ssRNA phage vOTUs (containing three core genes in any order). 11,222 vOTUs were assumed to represent near-complete genomes, given the presence of three full-length core genes [27].

### Characterisation of viral populations

All dsDNA vOTUs were annotated with Prokka v1.14.6 [68] using the Prokaryotic Virus Remote Homologous Groups (PHROGs) database [69] and the metagenome flag. Additional annotations were provided with eggNOG-mapper v2 [70, 71] with default settings. Genes putatively involved in metabolism were identified by clusters of orthologous groups (COGs): C, E, F, G, H, I, P and Q.

Taxonomic assessment of vOTUs was achieved with vConTACT2 v0.9.13 using “−rel-mode Diamond”, “−vcs-mode ClusterONE”, and a custom phage genome database (May 2020) [66] with all other settings set to default. The resultant genome network was visualised in R v4.0.5 using ggnet2 from GGally v2.1.2 [72] and the Fruchterman-Reingold force-directed algorithm. vOTUs were assigned into viral clusters (VCs) when clustering was significant (*p* < 0.05) and classified as outliers to the VC when clustering was non-significant. All unclustered vOTUs were classified as singletons.

ssRNA phages were classified into orders and families based on core protein isoforms [73], while genera and species were estimated using previously established RdRp gene clustering thresholds [27]. Phylogenetic assessment was performed on the concatenated core protein sequences aligned with MAFFT v7.271 [74]. Phylogenetic trees were constructed with FastTree v2.1.8 [75] using default settings and visualised in R using ggtree v2.5.3 [76–78].

Putative temperate phages were identified using previously described methods [79, 80]. Briefly, this identified temperate vOTUs encoding a protein associated with lysogeny or clustering with a known temperate phage. Additionally, vOTUs representing proviral sequences were assigned as temperate. Non-temperate vOTUs were assigned as lytic. Bacterial hosts were predicted using WIsH v1.0 [81] and a null model trained against 9,620 bacterial genomes as previously described [79]. Host predictions were filtered for *p* < 0.05 and were presented at the genus level.

### vOTU abundance and viral gene activity

DNA vOTU abundance was estimated by mapping DNA virome and total metagenome reads against a database of viral genomes (including non-redundant dsDNA vOTUs recovered in this study and all complete phage genomes in the custom phage database) using BBMap within BBTools [82] with “minid = 0.9”. vOTUs were only considered present in a sample if ≥ 75% of the contig length was covered ≥ 1 × by reads, as recommended [65, 83]. Given that the DNA virome and total metagenome libraries were only constructed in March 2017, DNA viral community compositions were only investigated at the stem extension growth stage. For detection of ssRNA phage vOTUs in RNA libraries, we used the above method and thresholds with the additional flag “ambig = random”. ssRNA phage community compositions were compared across seedling, stem extension, and pre-harvest growth stages. For the detection of DNA vOTU gene transcripts in RNA libraries, BAM files were sorted and indexed with SAMtools v1.10 [84]. BEDtools v2.26.0 [85] was used to extract read counts for each gene loci. Resulting read counts were filtered for ≥ 4 gene reads mapped across the replicates of each soil sample, with those < 4 converted to zero in each sample replicate. DNA vOTUs were identified as active when metatranscriptome reads mapped to ≥ 1 gene per 10 kb of the genome, as others have used previously [16].

### Data analysis and visualisation

All statistical analyses were conducted using R v4.0.5 [86]. Relative vOTU abundance values (counts per kilobase million, CPM) were computed by normalising read counts by genome length and library sequencing depth. The median of CPM values derived from the DNA virome and total metagenome libraries were computed to generate one abundance value per vOTU per sample. Viral community alpha diversity was described with Shannon’s *H* index computed on vOTU CPM profiles with phyloseq v1.34.0 [87]. Viral community beta diversity was described by computing a Bray-Curtis dissimilarity matrix from square-root transformed vOTU CPM profiles using vegan v2.5-7 [88], and subsequently visualised with non-metric multidimensional scaling (NMDS) ordination using vegan. Similarly, relative gene abundance values (transcripts per kilobase million, TPM) were computed by normalising read counts by gene length and library sequencing depth. Beta diversity in viral community activity was described in the same way as viral community composition. Two-way analysis of variance (ANOVA) tests and Tukey’s honestly significant differences (HSDs) were computed with stats v4.05. Permutational multivariate analysis of variance (PERMANOVA) tests and Mantel tests using Pearson’s product-moment correlation were performed with vegan. Linear mixed effect models were implemented using lmerTest v3.1-3 [89]. Differential abundance analysis was performed on raw read counts with DESeq2 v1.30.1 [90]. Plots were generated with ggplot2 v3.3.3 [91].

## Results

### Significant expansion of plant root-associated viruses identified from field-grown oilseed rape

To determine viral community composition across root/soil compartments and crop rotation practices, we recovered vOTUs from samples outlined in **Fig. 1**. A total of 1,059 non-redundant dsDNA vOTUs were recovered from the size-fractionated metagenome (DNA virome), total metagenome, and metatranscriptome libraries, with only one vOTU belonging to a previously isolated phage species (**Table S2**). The reconstruction of viral sequences from the metatranscriptome yielded 521 (49.8% of total) dsDNA vOTUs, which were not assembled from either of the DNA libraries (i.e., DNA virome and total metagenome). Additionally, a total of 16,541 non-redundant ssRNA phage vOTUs were recovered from the metatranscriptome, with 11,222 of these vOTUs representing near-complete ssRNA phage genomes.

**Fig. 1:**
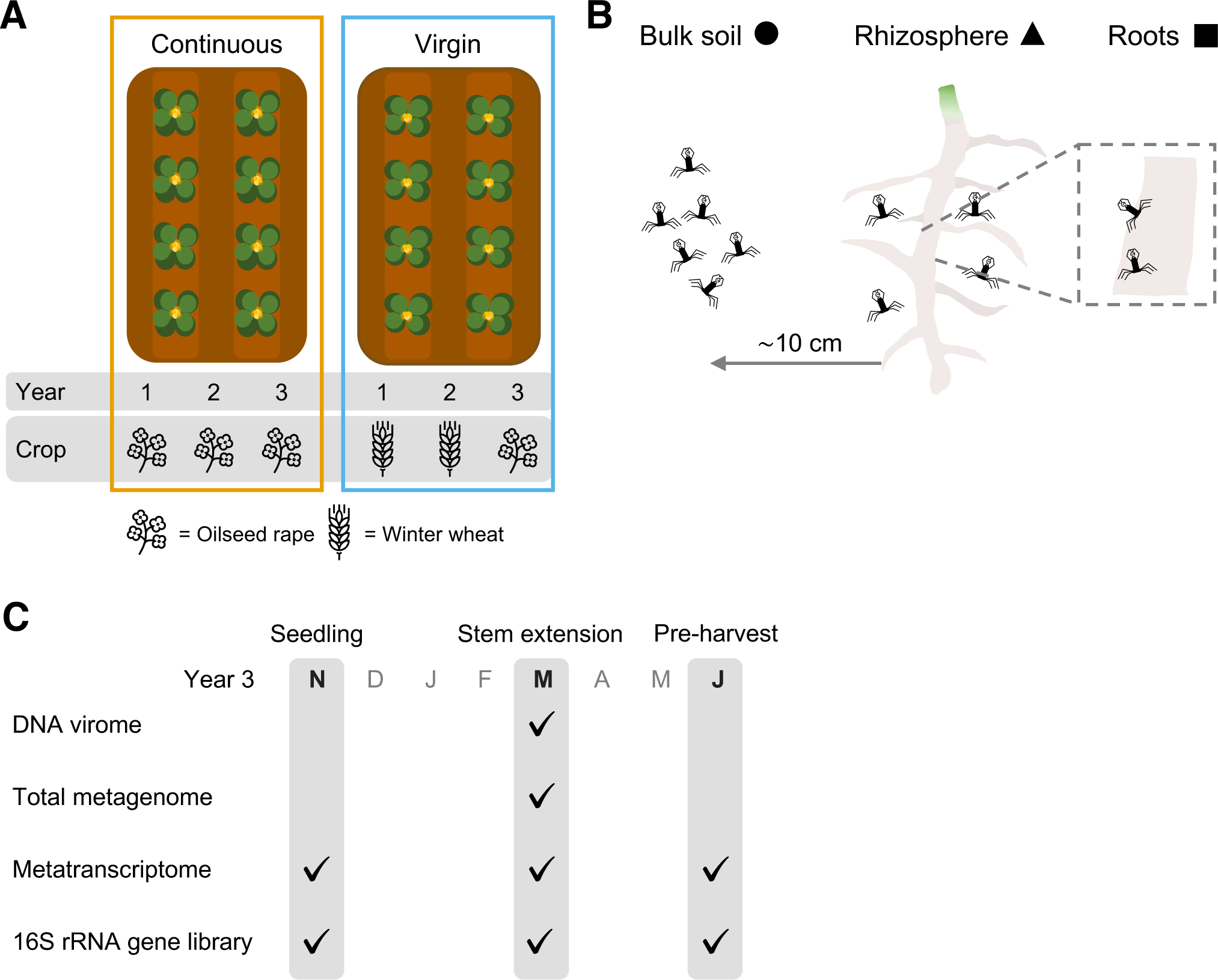
Overview of sampling strategy. **A** Schematic of sampled crop rotation practices. Four plots employed continuous cropping (left, orange) whereby oilseed rape was grown for three consecutive years. Another four plots employed virgin rotation (right, blue) whereby oilseed rape was grown following two preceding years of winter wheat. Eight plants were taken from each plot to generate each sample during the third year of crop growth. **B** Schematic of sampled compartments. Samples were taken from bulk soil (circles), rhizosphere soil (triangles), and roots (squares). **C** Libraries constructed for each sampling time point. Tick icon indicates data library construction for the given time point. Seedling samples were taken in November (N); stem extension samples were taken in March (M); pre-harvest samples were taken in June (J).

Next, we performed shared protein-based classification to investigate the similarity of recovered vOTUs with all currently available phage genomes, using vConTACT2 [92]. The resultant network contained viral clusters (VCs) representing roughly genus-level taxonomic groups (**Fig. 2A**); 262 (24.7% of total) dsDNA vOTUs and 7,677 (46.4% of total) ssRNA phage vOTUs formed 95 and 884 VCs, respectively (**Table S2; Table S3**). However, only 10 of these VCs contained phage genomes that had been previously isolated, demonstrating the undiscovered viral diversity found in this study. The proportion of dsDNA vOTUs forming genus-level VCs was similar across each library used to assemble vOTUs, and consistently lower than ssRNA phage vOTUs (**Fig. 2B**). Using previously established criteria [27], ssRNA phage vOTUs were resolved into 909 genera and 2,440 species within the class *Leviviricetes* (**Table S3**). This included 683 (75.1% of total) new genera and 2,379 (97.5% of total) new species, further highlighting the vast novel taxonomic diversity in the ssRNA phage vOTUs.

**Fig. 2:**
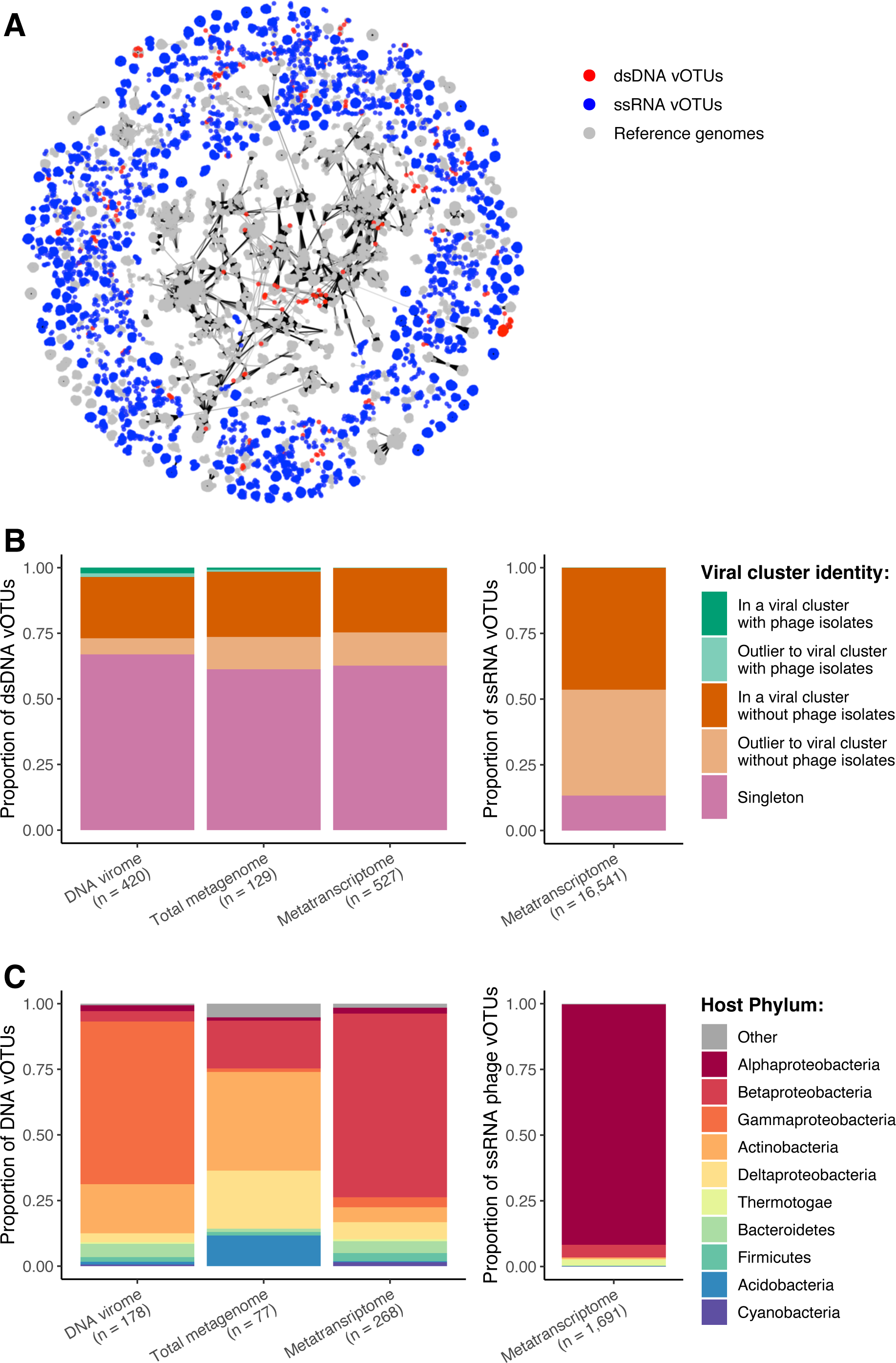
Taxonomic diversity and predicted hosts for recovered viral populations. **A** Shared protein content of recovered vOTUs with all currently available phage genomes, as determined by vConTACT2. Network graph visualisation includes 262 clustered dsDNA vOTUs (red nodes), 7,677 clustered ssRNA phage vOTUs (blue nodes), and 12,586 clustered reference DNA and RNA phage genomes (grey nodes). **B** Formation of genus-level viral clusters by recovery library. Relative proportion of vOTUs that formed viral clusters with previously isolated phage genomes (green), without previously isolated phage genomes (orange), and singletons i.e., those that are did not form viral clusters (pink) for dsDNA vOTUs (left) and ssRNA phage vOTUs (right). **C** Putative bacterial host phyla of vOTUs by recovery library. Relative proportion of vOTUs predicted to infect bacterial host phyla for dsDNA vOTUs (left) and ssRNA phage vOTUs (right). vOTUs with unknown host genera are excluded. Bar fill colour indicates bacterial host phylum for the top 10 most common host phyla. Proteobacteria are separated into classes. “Other” represents remaining host phyla.

Novel ssRNA phage diversity was further interrogated by constructing a phylogeny of 11,222 near-complete ssRNA phage vOTUs and all currently available *Leviviricetes* genomes (**Fig. S2**). 6,217 (55.4% of total) near-complete ssRNA phage vOTUs were resolved into 557 new genera, across all five existing *Leviviricetes* families (**Table S3**). This revealed the extension on existing *Leviviricetes* diversity found in other ecosystems [27–29, 31] and the expansion of the known number of *Leviviricetes* genomes by > 5 times.

To understand the potential ecological roles of soil viruses, we predicted the lifestyles and hosts of recovered vOTUs. Only 105 (9.9% of total) dsDNA vOTUs were predicted to represent temperate phages, indicating that the majority were likely to be obligately lytic (**Table S2**). In contrast, it was assumed that none of the ssRNA phage vOTUs were temperate, given that there has been no reported lysogeny among *Leviviricetes* phages. Bacterial hosts were predicted *de novo* for 518 (48.9% of total) dsDNA vOTUs and 1,691 (10.2% of total) ssRNA phage vOTUs using a probabilistic model [81]. The most common host taxonomic class varied depending on the library that the vOTUs were assembled from (**Table S2; Table S3**); gammaproteobacterial hosts were the most common among DNA virome-assembled vOTUs (61.2% of assigned hosts), actinobacterial hosts were the most common among total metagenome-assembled vOTUs (37.7% of assigned hosts), and betaproteobacterial hosts were the most common among metatranscriptome-assembled vOTUs (68.7% of assigned hosts) (**Fig. 2C**). A single, uncultured alphaproteobacterial host genus was the most common among ssRNA phage vOTUs (91.5% of assigned hosts) (**Fig. 2C**). 68/85 (80.0%) of the bacterial genera putatively infected by dsDNA vOTUs were detected in our soil samples, while only 3/12 (25.0%) of the bacterial genera putatively infected by ssRNA phage vOTUs were detected (**Table S4**).

The prevalence of the vOTUs recovered in this study was compared by detecting vOTU sequences through read mapping, from the DNA libraries (for dsDNA vOTUs) and metatranscriptomes (for ssRNA vOTUs). Despite sampling not achieving a richness asymptote, 698 DNA vOTUs were detected in at least one sample (**Fig. S3A**). This included 382/420 (91.0%) DNA virome-assembled vOTUs, 116/129 (89.9%) total metagenome-assembled vOTUs, 215/527 (40.8%) metatranscriptome-assembled vOTUs, and one previously isolated ssDNA phage genome (**Table. S2**). DNA virome-assembled vOTUs were detected in a mean of 1.60 samples and represented 80.9% of all DNA vOTUs present in only one sample. Total metagenome-assembled vOTUs were detected in a mean of 2.83 samples, while metatranscriptome-assembled vOTUs were detected in a mean of 4.31 samples and represented 70.6% of the vOTUs detected in at least half of the samples. As with the dsDNA vOTUs, the sampling of ssRNA phage vOTUs did not reach a richness asymptote, with 12,162/16,541 (73.5%) vOTUs detected in at least one sample metatranscriptome (**Fig. S3B**).

By mapping metatranscriptome reads to vOTU gene sequences, we identified 827 (78.1% of total) active dsDNA vOTUs, including 296/420 (70.5%) DNA virome-assembled vOTUs, 104/129 (80.6%) total metagenome-assembled vOTUs, and 444/527 (84.3%) metatranscriptome-assembled vOTUs (**Table S5**). Additionally, 63 previously isolated dsDNA and ssDNA phage genomes were identified as active in at least one sample. The median relative activity of dsDNA vOTUs assembled from the metatranscriptome was 10.2 and 11.6 times greater than vOTUs assembled from the total metagenome and DNA virome, respectively.

### Plant root association and crop rotation shapes both DNA and RNA viral community composition

compartment in driving the diversity of both DNA viral communities (*F* = 5.116, *df* = 1, *p* = 0.0431) and ssRNA phage communities (*F* = 101.344, *df* = 2, *p* < 0.0001). For subsequent analyses of ssRNA phage communities across compartments, we chose to exclude the roots given that their very low richness and diversity inflated overall compartmental differences.

Next, we investigated the dissimilarities between viral communities through NMDS ordinations (**Fig. 3B**). PERMANOVA tests identified that both compartment (8.6-14.7% variance) and crop rotation (10.3-19.2% variance) had significant contributions to the differences in viral community composition at each growth stage (**Fig. 3C; Table S6**). Despite compartment contributing to > 2 times the variation in co-existing bacterial community composition, the contribution of crop rotation was similar for viruses and bacteria. Subsequently, Mantel tests revealed significant correlations between bacterial community composition and both DNA viral communities (*r* = 0.3301, *p* = 0.0039) and ssRNA phage communities (*r* = 0.4642, *p* = 0.0001).

**Fig. 3:**
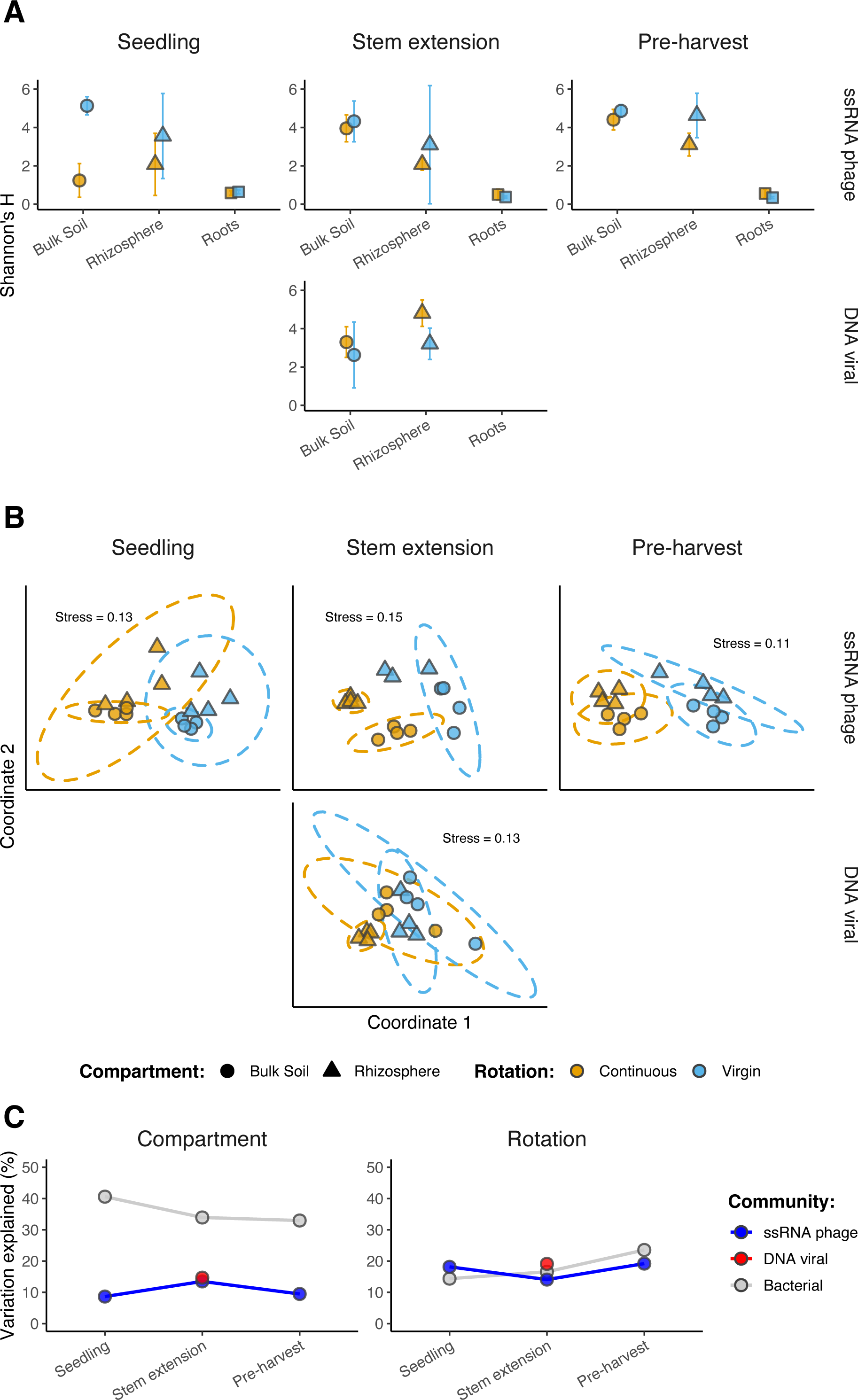
Diversity in viral community composition. **A** Alpha diversity of DNA viral community composition and ssRNA phage community composition. Mean alpha diversity indexes (Shannon’s *H*) for each viral community composition across compartments, at each crop growth stage. Shapes are coloured based on field crop rotation strategy: continuous cropping (orange) and virgin rotation (blue). Shapes indicate compartment: bulk soil (circles), rhizosphere soil (triangles), and roots (squares). Error bars denote a 95% confidence interval around the mean. **B** Beta diversity of DNA viral community composition and ssRNA phage community composition. Non-metric multidimensional scaling (NMDS) ordination plots, representing the dissimilarities between community compositions, for each growth stage. Ordinations represent community compositions containing ssRNA phages at seedling (n = 8,125), ssRNA phages at stem extension (n = 6,936), DNA viruses at stem extension (n = 698), and ssRNA phages at pre-harvest (n = 10,998). Shapes are coloured based on field crop rotation strategy: continuous cropping (orange) and virgin rotation (blue). Shapes indicate compartment: bulk soil (triangles) and rhizosphere soil (circles). Stress values associated with two-dimensional ordination are reported for each plot. **C** Variation in community composition explained by soil compartment and crop rotation. PERMANOVA results describe the variance in community composition explained by soil compartment and crop rotation, respectively, for each growth stage. Points are coloured based on community: ssRNA phage community (blue), DNA viral community (red), and bacterial community (grey).

Given the vast richness of *Leviviricetes* found in this study, we interrogated compartmental differences further by describing the composition of *Leviviricetes* families across root, rhizosphere soil, and bulk soil compartments (**Fig. S4**). This indicated a high degree of spatial structuring among ssRNA phage communities, even at the family level, with additional smaller effects of crop rotation and growth stage.

### Continuous cropping drives the emergence of distinct, active DNA viruses in seedling rhizospheres

To uncover the potential drivers of viral activity, we first explored the number of active DNA vOTUs detected in metatranscriptomes across root/soil compartments over time (**Fig. S5**). A two-way ANOVA test was performed, revealing significant effects of compartment (*F* = 218.546, *df* = 2, *p* < 0.0001), crop rotation (*F* = 139.185, *df* = 1, *p* < 0.0001), and growth stage (*F* = 508.088, *df* = 2, *p* < 0.0001) on active vOTU prevalence. It took until stem extension for differences between rotation practices to be observed in the bulk soil and roots, while differences in rhizosphere soil were apparent from the seedling stage (**Fig. S5**). In fact, there were 196 active vOTUs detected in the seedling rhizosphere under continuous cropping which were absent in the seedling rhizosphere under virgin rotation. The relative activity of these vOTUs increased over time (*F* = 51.764, *df* = 2, *p* < 0.0001), particularly in rhizosphere soil under continuous cropping (**Fig. S6**).

To investigate the ecological consequences of active vOTUs on their hosts, we trained linear mixed effect models, using compartment as a random effect. This revealed significant linear relationships between active vOTUs and both bacterial host abundance (*b* = −0.039, *p* < 0.0001, **Table S7**; **Fig. S7A**) and bacterial community alpha diversity (*b* = 0.001, *p* = 0.0147, **Table S8**; **Fig. S7B**).

### Rhizosphere enrichment of DNA viral activity displays a spatial gradient

The dissimilarity between total viral community activity at each growth stage was investigated with NMDS ordinations (**Fig. 4A**). PERMANOVA tests revealed the significant and dynamic contributions of both compartment (22.5-35.8% variance) and crop rotation (19.1-41.0% variance), such that the effect of crop rotation increased over time, while the effect of compartment decreased (**Fig. 4B; Table S9**). Comparing these effects on the activity of vOTUs assembled from each library independently revealed that compartmental differences were greatest among metatranscriptome-assembled vOTUs (**Fig. S8; Table S10**).

**Fig. 4:**
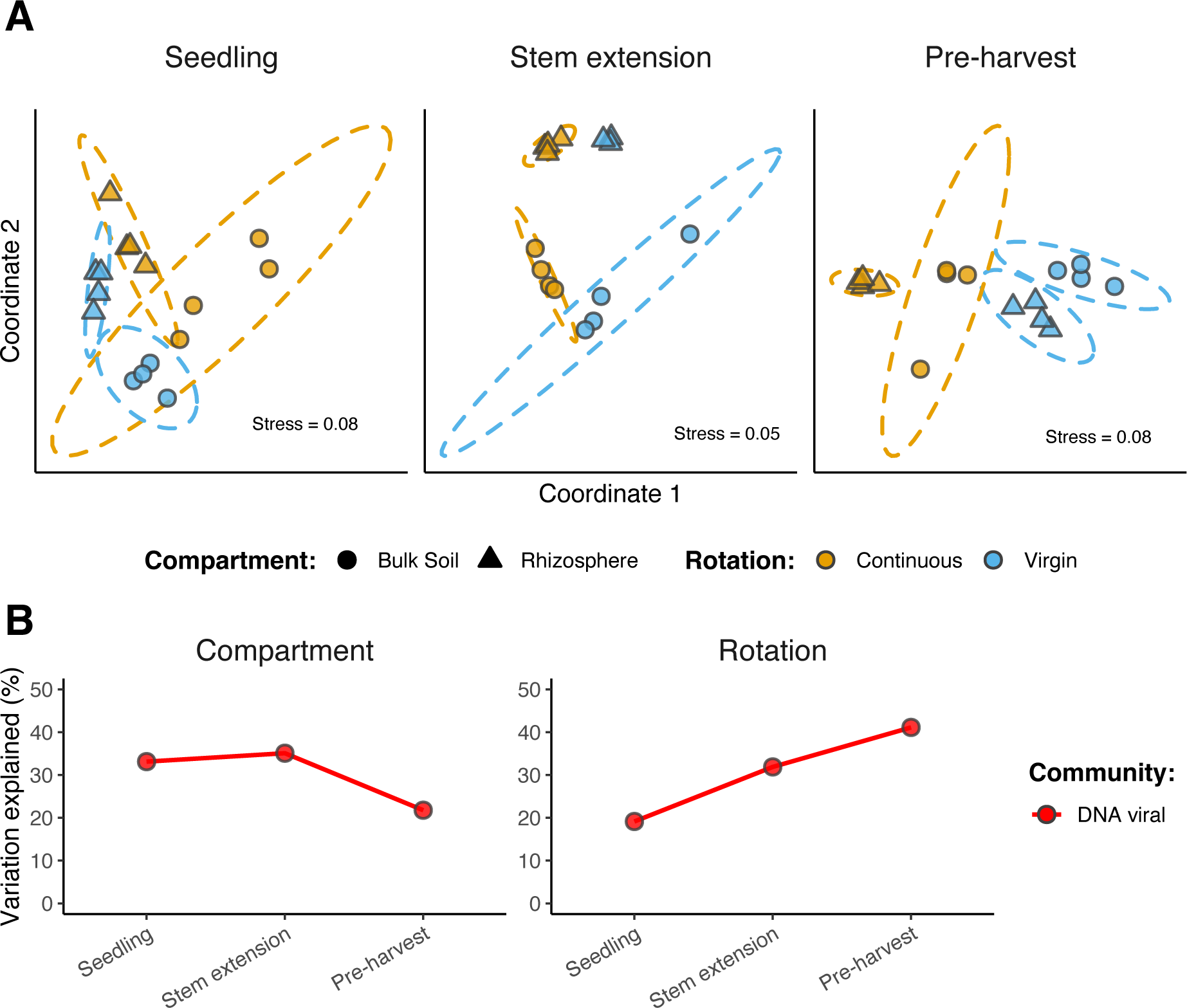
Diversity in viral community activity. **A** Beta diversity of DNA viral community activity. Non-metric multidimensional scaling (NMDS) ordination plots, representing the dissimilarities between gene transcript abundances, for each growth stage. Ordinations represent DNA viral activity at seedling (n = 6,696), DNA viral activity at stem extension (n = 7,958), and DNA viral activity at pre-harvest (n = 11,299). Shapes are coloured based on field crop rotation strategy: continuous cropping (orange) and virgin rotation (blue). Shapes indicate compartment: bulk soil (circles) and rhizosphere soil (triangles). Stress values associated with two-dimensional ordination are reported for each plot. **B** Variation in community activity explained by soil compartment and crop rotation. PERMANOVA results describe the variance in viral community activity explained by soil compartment and crop rotation, respectively, for each growth stage. Points are coloured based on community: DNA viral (red).

To further interrogate compartment-specific viral activity, we first identified differentially active viral genes in either rhizosphere soil or bulk soil. This found ∼ 14 times more genes (3,589 vs. 250) with significantly greater activity in the bulk soil (relative to rhizosphere soil) than in rhizosphere soil (relative to the bulk soil). We then compared the total activity of these genes across all compartments, revealing that > 78% of viral community activity was soil compartment enriched (**Fig. S9**). Furthermore, rhizosphere-enrichment of viral activity displayed a spatial gradient, representing a greater proportion of total community activity at the roots than in rhizosphere soil.

Given that previous research has associated viruses with modulating their hosts’ metabolism, we investigated the proportion of viral community activity encoding metabolic functions (**Fig. S10**). A two-way ANOVA was performed, identifying significant effects of both compartment (*F* = 67.682, *df* = 2, *p* < 0.0001) and growth stage (*F* = 11.098, *df* = 2, *p* < 0.0001) on viral-encoded metabolic activity.

## Discussion

### Novel, diverse, and active ssRNA phages in plant root-associated ecosystems

In the present study, we have demonstrated that ssRNA phages were both abundant and active across root, rhizosphere soil and bulk soil compartments. In doing so, we have expanded the number of *Leviviricetes* genomes by > 5 times and identified 683 new genera and 2,379 new species (**Table S3**). The existing phylogeny defined from a variety of ecosystems remained stable with the addition of the new viral sequences we recovered [27–29, 31, 73] (**Fig. S2**). In addition to uncovering novel diversity, we discovered compartmental differences in ssRNA phage communities (**Fig. 3C**), such that the composition of *Leviviricetes* families varied with proximity to crop roots (**Fig. S4**). This is the first time that ssRNA phage communities have been investigated at the root surface, and the first evidence of plant roots shaping their community composition. In combination with recent investigations of *Leviviricetes* in terrestrial ecosystems [30, 31], our discovery emphasises the underappreciation of RNA viruses in soils, relative to DNA viruses.

We hypothesise that the trend of *Leviviricetes* across compartments mirrors host populations, given the significant correlation observed between phage and host communities, and that phages require their hosts to replicate. This phenomenon is indicative of predator-prey dynamics, which have been previously demonstrated to link phage and bacterial population abundances in soil crust [93]. Furthermore, the dynamic changes in the relative abundances of ssRNA phages is likely to represent substantial viral reproduction, indicating the active infection of bacterial hosts. ssRNA phages have been established to infect *Pseudomonadota* [94, 95], a highly abundant soil phylum including key participants in the cycling of carbon, nitrogen, and sulphur [96]. Therefore, by driving the turnover of bacteria, the diverse, abundant, and active ssRNA phages recovered in this study are expected to impact terrestrial biogeochemical cycling. Our *de novo* host predictions suggest that ssRNA phages infect additional bacteria outside of the current *Alphaproteobacteria* and *Gammaproteobacteria* model systems [94, 95] (**Fig. 2C**). This highlights the underestimated potential of ssRNA phages in the turnover of host populations across root-associated ecosystems. Thus, the development of further model systems, involving cultivation of both phage and its host, is imperative to investigate the impacts of ssRNA phages on rhizosphere ecology.

### Viral-host interactions across root-associated microbiomes

Through their detection across root, rhizosphere soil, and bulk soil compartments, we were able to implicate active DNA viruses in shaping co-existing bacterial communities. The majority of the vOTUs recovered in this study were likely to represent lytic dsDNA phages, given almost half of the dsDNA vOTUs had predicted bacterial hosts (**Fig. 2C**) and the prevalence of lysogeny was low (**Table S2**). Lytic viral activity was evidenced by a negative association between active vOTUs and co-existing bacterial host abundance (**Figure S7A**; **Table S7**). Furthermore, active vOTU prevalence was positively associated with bacterial community diversity (**Figure S7B**; **Table S8**), potentially implicating viral activity in driving bacterial diversity (or vice versa). These are features of the “Kill-the-Winner” hypothesis, which predicts that the population growth of dominant bacterial species is limited by viral lysis [97, 98]. In driving the turnover of host populations, phages are likely to contribute to bacterial community succession and the maintenance of community diversity. Despite there being very limited previous evidence of Kill-the-Winner dynamics occurring in soils [93], other predator-prey dynamics, notably bacterial predation by protists and nematodes, has long been associated with changes to microbial community composition [99]. But given that viruses are more selective in their infection of specific host taxa, viral-host interactions are likely to be more variable in space and time. Accordingly, the increasing prevalence of active vOTUs observed across the growing season (**Fig. S5**) suggests that Kill-the-Winner dynamics exhibits temporal variation in soil. This implies that the impact of soil viral activity may increase over the crop growth season, as the root-associated microbiome matures.

By comparing the activity of viral communities between soil compartments, we observed that most viral activity was enriched in either rhizosphere soil or bulk soil (**Fig. S9**). Moreover, we revealed a spatial gradient of viral activity across the root-associated microbiomes, indicating that viruses could have soil niche-specific functions. Viral infection can result in the modulation of microbial host metabolism through the expression of viral-encoded AMGs, as evidenced in marine ecosystems [4–7]. The previous identification of AMGs relating to carbon acquisition and processing has implicated soil viruses in terrestrial carbon and nutrient cycling [16, 39, 42, 43]. Here, we have extended these previous efforts by characterising the activity of viral-encoded metabolic genes, whose combined relative activity increased with proximity to crop roots (**Figure S10**). This parallels the enrichment of microbial activity in the root-associated microbiomes [35]. Thus, soil viruses may contribute to rhizosphere ecology and function through the augmented reprogramming of host metabolism, which could act either antagonistically or synergistically with plants in the control of their root-associated microbiomes [54].

### Viral priming in the crop rhizosphere

Previous studies have demonstrated the impacts of crop management practices on bacterial, fungal, and nematode communities in the rhizosphere [49, 50]. By comparing viral communities associated with continuous cropping and virgin rotation, we have provided the first evidence of crop rotation-driven impacts on soil microbial communities extending to viruses. In fact, differences in viral community composition (**Fig. 3C**) and activity (**Fig. 4B**) were sometimes greater between crop rotation practices than between soil compartments. Furthermore, we detected significantly more active vOTUs in the seedling rhizosphere under continuous cropping than in the seedling rhizosphere of the crop grown in virgin rotation (**Fig. S5**). We propose the principal of “viral priming” to explain this observation; viruses remaining in the soil from the previous growing season were adapted to infect hosts colonising the juvenile rhizosphere in the current season, only under continuous cropping. Put simply, a greater number of viruses were actively infecting hosts to which they had been previously exposed to. Subsequent differences between rotation practices at the seedling stage were limited to the rhizosphere given the similarity of bacterial communities observed in the bulk soil (**Fig. S11**). The required persistence of viruses between growth seasons could have been facilitated by (i) the continuous presence of susceptible hosts, (ii) clay particles providing protection from degradation [100], and/or (iii) low soil temperatures preventing viral inactivation [101].

The local adaptation that allows priming viruses to infect hosts colonising the rhizosphere can be explained by antagonistic coevolution. A study in soil microcosms demonstrated that the fitness cost of phage resistance among bacteria limited their resistance to include only co-occurring phages [102]. Meanwhile among soil phages, a high level of local adaptation has been shown to result in greater infection rates of co-existing rhizobia strains, as compared to geographically distant strains [103]. Therefore, given the elevated fitness cost of resistance, specific to the rhizosphere [54], newly colonising bacteria are likely to be susceptible to primed viruses. Patterns of phage-bacteria coevolution have previously been observed on the centimetre scale within soils [104], indicating the feasibility for viral priming to occur specifically in the rhizosphere. In contrast, under virgin rotation, viruses remaining in the soil following the harvest of wheat were maladapted to infect the distinct bacterial community that colonised the seedling rhizosphere of the new crop (**Fig. S11**). Continuous crop growth has been used to explain the accumulation of plant fungal pathogens in rhizosphere soil, which were shown to result in crop yield decline [51]. We speculate that greater viral activity under continuous cropping, due to viral priming, could play a role in regulating both deleterious and beneficial plant-microbe interactions, thus impacting plant health and yield. Moreover, given that the activity of primed viruses increased across growth stages (**Fig. S6**), there is likely to be a significant and increasing impact of viral priming on the root-associated microbiomes throughout the growing season. While the net positive or negative consequences of viral priming are yet to be elucidated, we have provided evidence that crop rotation mitigates viral priming activity in the rhizosphere.

### Combining metatranscriptomics with metagenomics and viromics to study soil viral communities

We also demonstrate that integrating metatranscriptomics with conventional DNA-based omics approaches mitigates any potential failure to capture ecologically significant viral communities. To describe viral populations, we simultaneously recovered viral genomes from a DNA virome, total metagenome, and metatranscriptome. Remarkably, almost half of the dsDNA vOTUs presented here were assembled from the metatranscriptome alone (**Table S2**), despite there being no precedent for this recovery method in previous viral ecology studies. Different vOTUs can be recovered between DNA libraries as a result of the library preparation method used, particularly size-filtration to obtain the DNA virome, which has been confirmed to underrepresent viruses with larger capsid sizes [105]. However, this is the first time that DNA viral genomes have been simultaneously recovered from DNA and RNA libraries using the same viral prediction tools and thresholds.

The average prevalence of metatranscriptome-assembled vOTUs was greater than those assembled from DNA libraries, which may have been responsible for our ability to observe greater compartmental differences among these vOTUs (**Fig. S8**). Previously, total metagenomes have been shown to bias towards the most persistent viruses, capable of infecting the most abundant host organisms [106]. However, many of the highly prevalent metatranscriptome-assembled vOTUs eluded recovery from the total metagenome. Recently it has become apparent that the hyper-modification of phage DNA prevents the sequencing of certain phage genomes [107, 108]. Subsequently, many phage genomes remain absent in DNA metagenomic samples prepared using transposon-based library methods, as used in this study. Transcriptomics has previously been used to assemble the genome of phage YerA41 from phage-infected cells, thus overcoming the unknown DNA modification that prevented DNA sequencing [109]. The assembly of phage genomes eluding standard DNA sequencing methods, in addition to differences in library sizes, could explain why so many dsDNA vOTUs were exclusively recovered from the metatranscriptome.

Upon further investigation of the vOTUs assembled from each library, we observed consistently high taxonomic novelty (**Fig. 2B**), but large shifts in the most common bacterial host phyla (**Fig. 2C**). This indicates possible ecological differences in the viruses accessed by each method, highlighting the value of their combination for describing viral ecology. In addition to its role in reconstructing viral genomes, we implemented metatranscriptomics to detect active vOTUs and characterise viral community activity. Subsequently, we were able to distinguish the ecologically active viral fraction from the “banked” viruses remaining dormant in viral communities at the time of sampling [15, 110]. Furthermore, we have demonstrated that the metatranscriptome accessed the most active viruses, which are vital for investigations of viral ecology, given that viral activity implies the presence and susceptibility of co-existing host organisms. In fact, almost half of the rhizosphere priming viruses were exclusively accessed by the metatranscriptome. This presents the metatranscriptome as a useful, yet underutilised, tool to study soil viral communities.

## Conclusions

In summary, we aimed to investigate the influence of crop rotation on the composition and activity of bulk soil, rhizosphere soil, and root viral communities. Combining viromics, metagenomics, and metatranscriptomics, we recovered 1,059 dsDNA vOTUs, with almost half of them assembled from the metatranscriptome alone. We also recovered thousands of ssRNA phage vOTUs, including 683 new genera and 2,379 new species, and expanding the number of *Leviviricetes* genomes by > 5 times. By describing ssRNA phage communities at the root surface for the first time, we emphasise their underappreciation in soil, as compared to DNA viruses. Furthermore, we revealed spatiotemporal viral activity indicative of “Kill-the-Winner” dynamics and postulate that viral reprogramming of host metabolism is greater in the rhizosphere than in bulk soil. We also provided the first evidence of crop rotation-driven impacts on soil microbial communities extending to viruses, proposing the novel principal of “viral priming” in the rhizosphere. Our work demonstrates that the roles of soil viruses need greater consideration to exploit the rhizosphere microbiome for food security, food safety, and environmental sustainability. Future studies should continue to investigate soil viral activity with relation to rhizosphere ecology to provide a framework by which we can manage viral communities within agricultural ecosystems. Critically, viruses should be universally included in plant microbiome studies, particularly where these microbiomes have implications for agricultural productivity.

## Abbreviations

AMG: Auxiliary Metabolic Gene
ANI: Average Nucleotide Identity
ANOVA: ANalysis Of Variances
Bp: base pairs
CPM: Counts Per Million
dsDNA: double-stranded DNA
HSD: Honestly Significant Differences
Kb: kilobases
NMDS: Non-metric Multi-Dimensional Scaling
OTU: Operational Taxonomic Unit
PERMANOVA: PERmutational Multivariate ANalysis Of Variances
ssRNA: single-stranded RNA
VC: Viral Cluster
vOTU: Viral Operational Taxonomic Unit.

## Acknowledgements

We acknowledge the use of MRC-CLIMB for the provision of high-performance servers, without which this work wouldn’t be possible. We would also like to thank Howard Hilton for kindly providing the photograph used in Figure S1.

## Declarations

### Ethics approval and consent to participate

Not applicable.

### Consent for publication

Not applicable.

### Availability of data and materials

Post-QC reads are available from the European Nucleotide Archive (ENA) under the Study Accession PRJEB49313. Sample Accession information is included in Table S1. dsDNA vOTU genome sequences were deposited to ENA under Sample Accession SAMEA12363644. ssRNA phage vOTU genome sequences were deposited to ENA under Sample Accession SAMEA11777518. FASTA nucleotide files containing vOTU genomes, FASTA amino acid files containing vOTU genes, vOTU gene annotations, FASTA amino acid files containing ssRNA phage core protein sequences, Newick tree file containing ssRNA phage phylogeny, and vConTACT2 network input and output files are available from figshare (https://figshare.com/XXX). The custom R script used to generate figures and tables, along with required input files, are available from GitHub (https://github.com/GeorgeMuscatt/ RhizosphereVirome).

### Competing interests

The authors declare that they have no competing interests.

### Funding

G.M. was funded by the EPSRC & BBSRC Centre for Doctoral Training in Synthetic Biology grant EP/L016494/1. A.M. was funded by MRC grants MR/L015080/1 and MR/T030062/1. G.B. was funded by BBSRC grant BB/L025892/1. E.J. was funded by Warwick Integrative Synthetic Biology (WISB), supported jointly by BBSRC & EPSRC, grant BB/M017982/1.

### Authors’ contributions

E.M.H.W., C.Q., A.M., G.D.B. and E.J. conceived and designed the experiment. G.T. and G.D.B. designed and managed the field experiment. S.H. and I.D.E.A.L. collected samples and extracted nucleic acids. G.M. and S.R. performed read processing and preparation. G.M. carried out bioinformatic analyses, generated R scripts, interpreted data, prepared figures, and produced the first draft of the manuscript. I.D.E.A.L., A.M., G.D.B., and E.J. provided edits and additional contributions to the manuscript. All authors read and approved the final submitted manuscript.

---

**Supplementary Figure S1:**
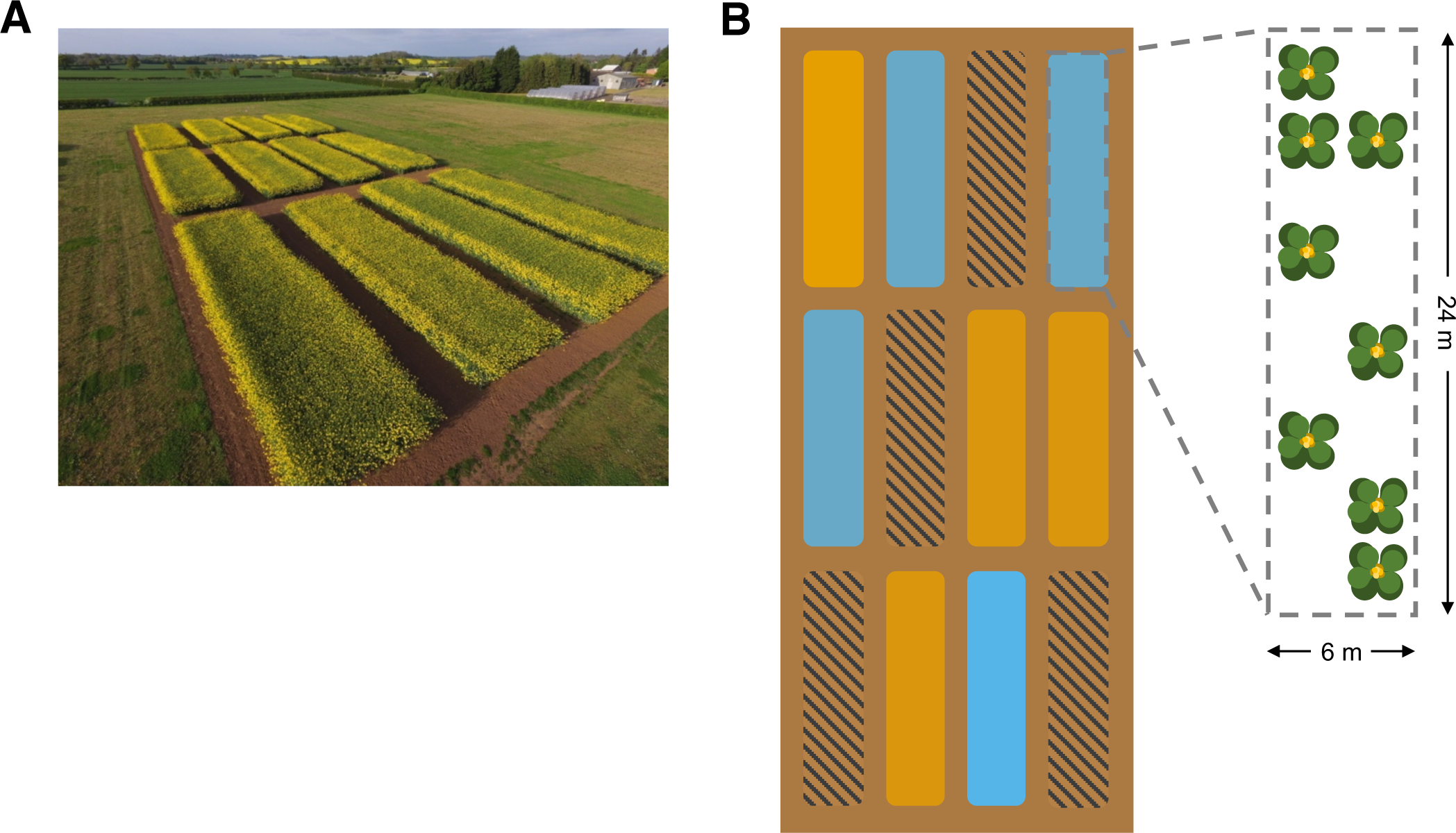
Overview of field site. **A** Photograph of field site, including twelve plots, taken between stem extension and pre-harvest growth stages in the third year of the trial in 2017. **B** Schematic of field site, representing twelve plots. Coloured plots indicate the two crop rotation practices sampled in this study: continuous cropping (orange) and virgin rotation (blue). Hashed plots were not sampled in this study. Expanded plot indicates that eight plants were sampled per plot to represent one replicate of each crop rotation practice.

**Supplementary Figure S2:**
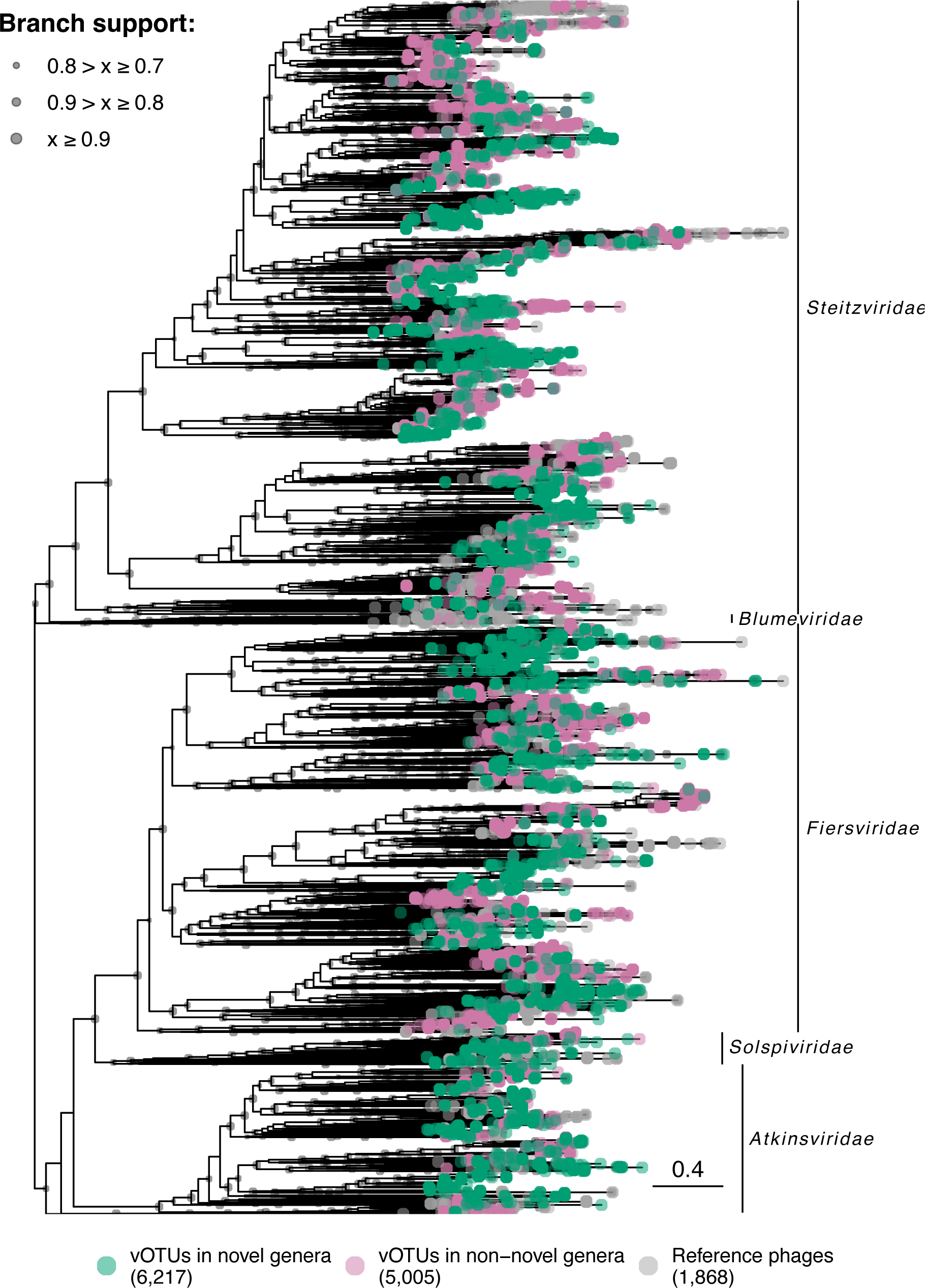
Phylogenetic assessment of ssRNA phage vOTUs. Phylogeny of ssRNA phage vOTUs using concatenated core protein sequences (maturation protein, coat protein, and RNA-dependent RNA polymerase). Phylogenetic tree contains 1,868 existing *Leviviricetes* phage sequences and our 11,222 full-length ssRNA phage vOTUs. Branch tip colours indicate novelty of genome sequence: vOTUs in novel genera (green, n = 6,217), vOTUs in non-novel genera (pink, n = 5,005), and existing ssRNA phage genomes (grey, n = 1,868). Clade labels indicate current *Leviviricetes* families. Brand node labels indicate branch support: ≥ 0.9 (large circles), ≥ 0.8 (medium circles), ≥ 0.7 (small circles), < 0.7 (no circle).

**Supplementary Figure S3:**
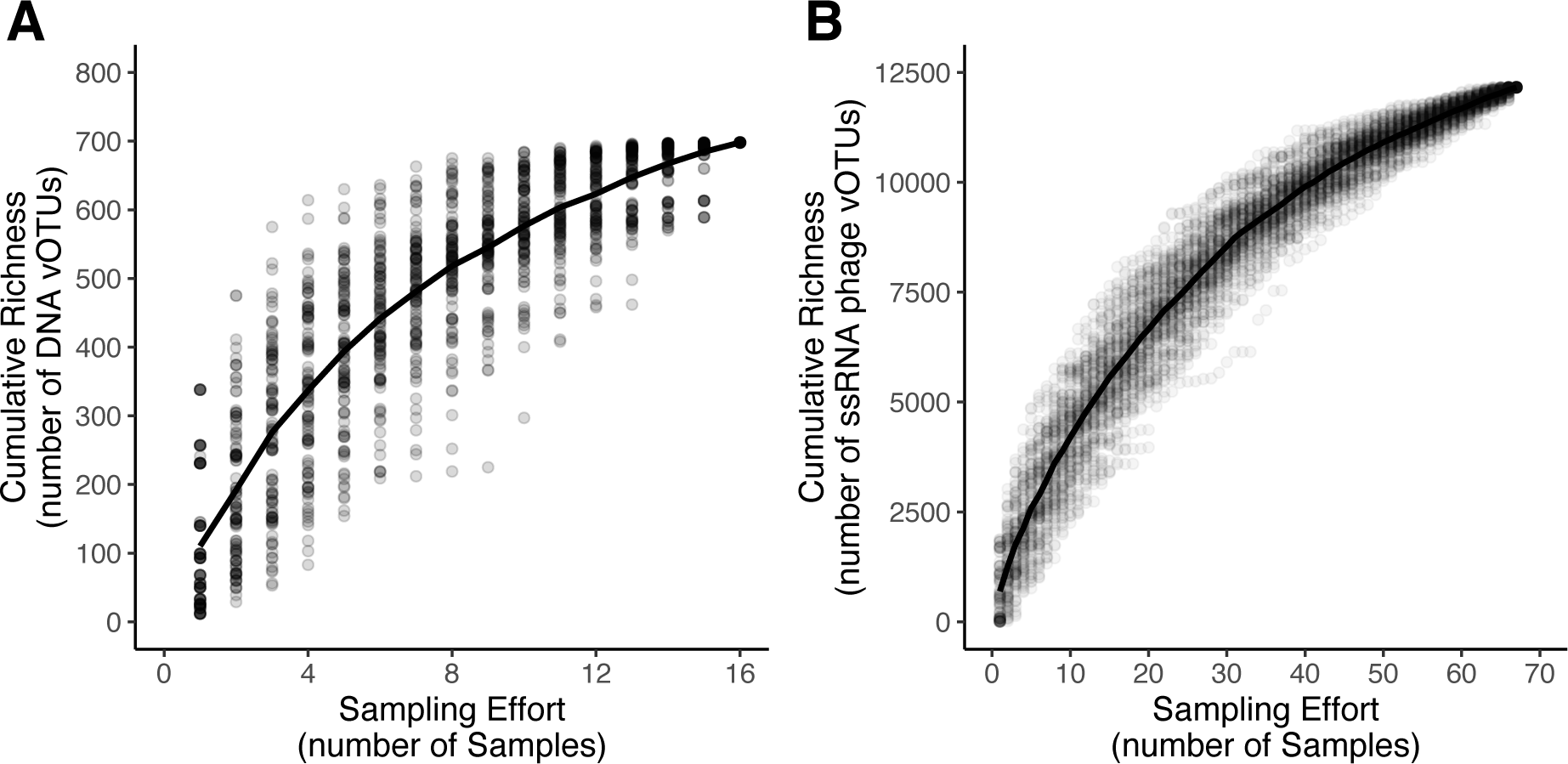
Accumulation curves of vOTUs. Accumulation curves for **A** DNA vOTUs and **B** ssRNA phage vOTUs. Dots represent 100 permutations of cumulative richness at each sampling effort. Line indicates the mean cumulative richness in vOTUs detected.

**Supplementary Figure S4:**
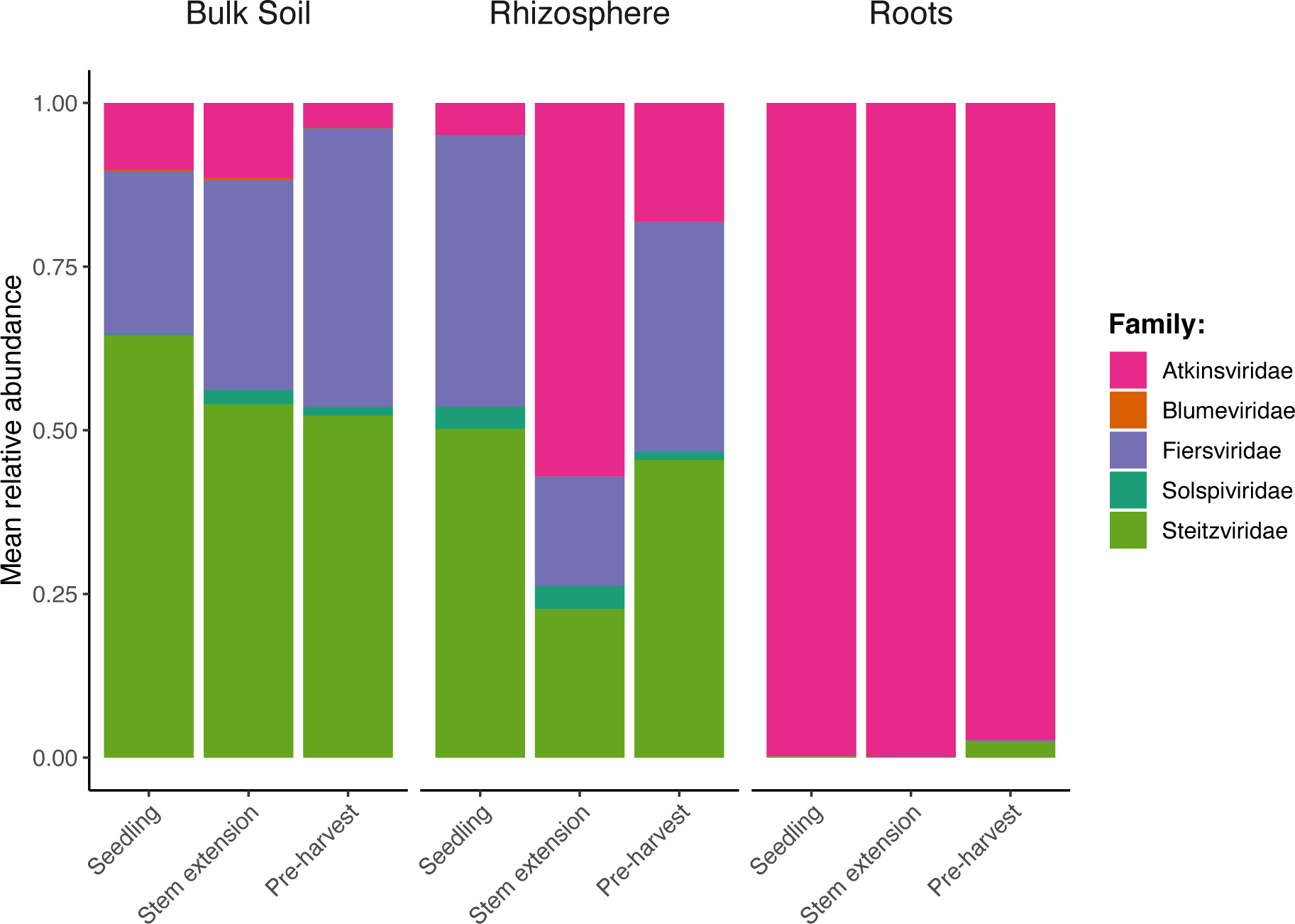
Summed mean relative abundance of *Leviviricetes* families. Relative abundance of *Leviviricetes* families in each root/soil compartment, across growth stages. Colour indicates *Leviviricetes* family: *Atkinsviridae* (pink), *Blumeviridae* (orange), *Fiersviridae* (purple), *Solspiviridae* (blue-green), and *Steitzviridae* (green).

**Supplementary Figure S5:**
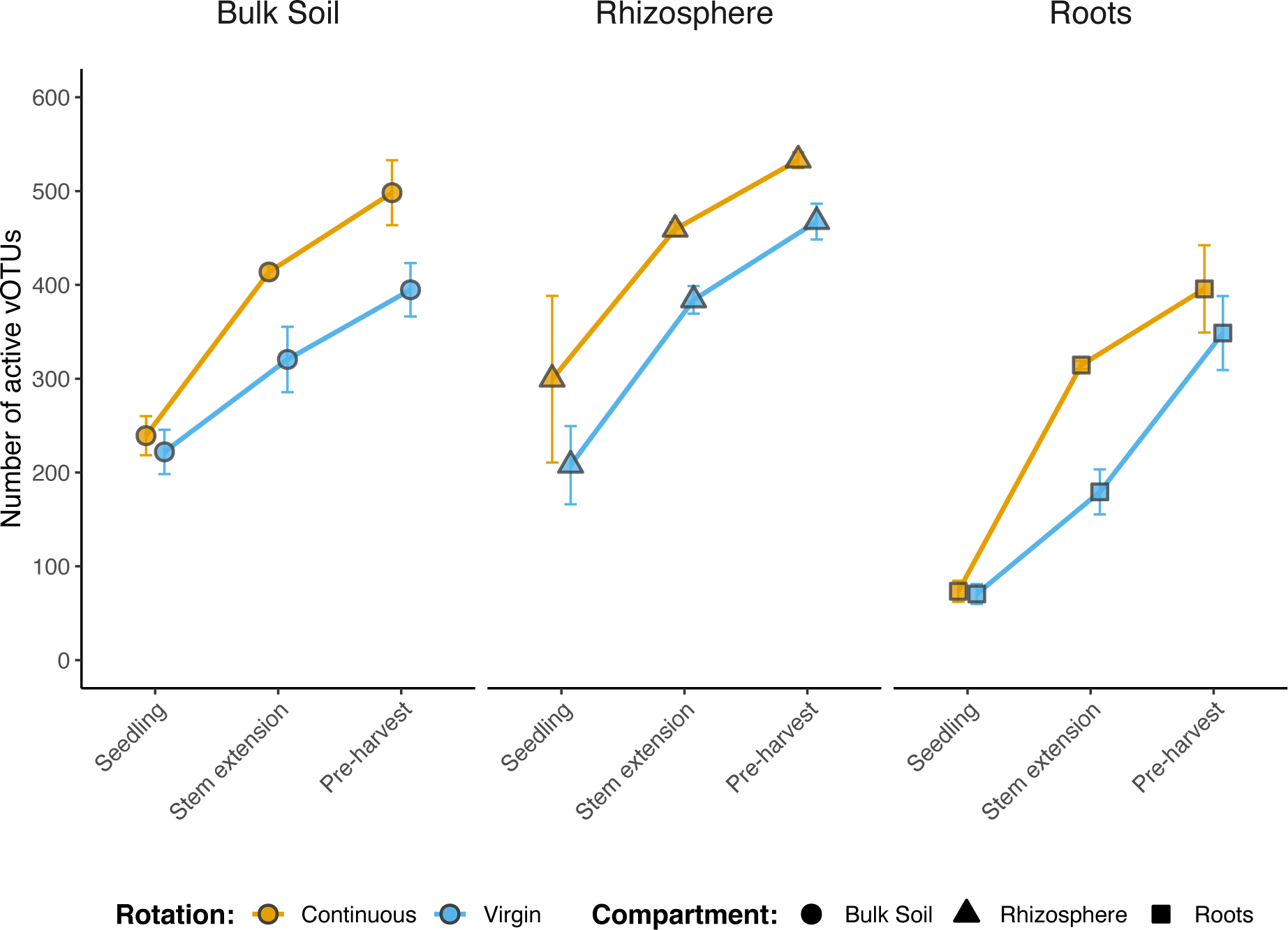
Detection of active vOTUs. Mean number of active DNA vOTUs detected in each root/soil compartment, across growth stages. Shapes are coloured based on field crop rotation strategy: continuous cropping (orange), virgin rotation (blue). Shapes indicate compartment: bulk soil (circles), rhizosphere soil (triangles), and roots (squares). Error bars denote a 95% confidence interval around the mean.

**Supplementary Figure S6:**
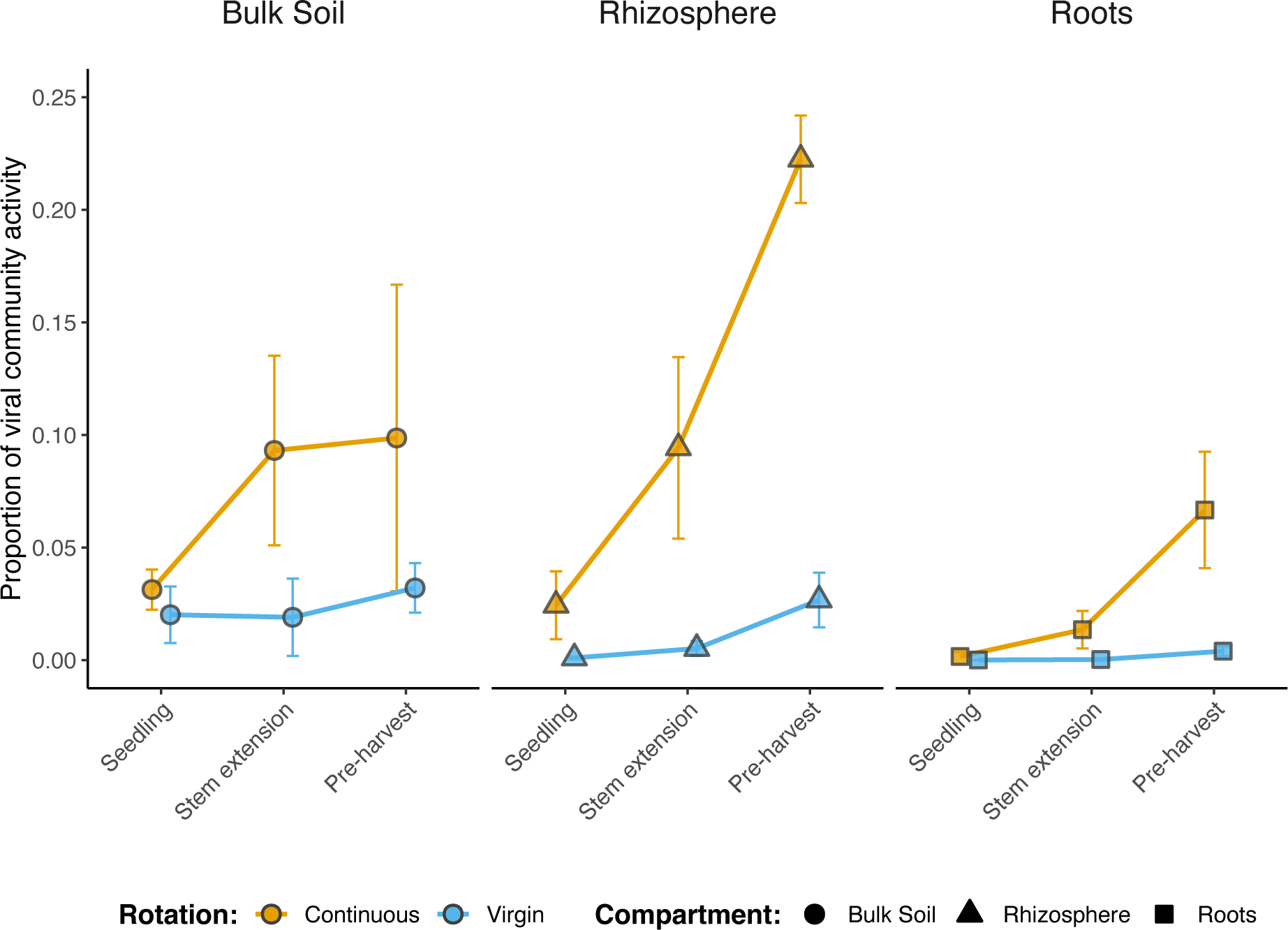
Summed mean relative activity of rhizosphere-priming vOTUs. Mean proportion of viral community activity represented by rhizosphere-priming vOTUs (n = 196) in each root/soil compartment, across growth stages. Rhizosphere-priming vOTUs were detected in the seedling rhizosphere under continuous cropping but were absent in the seedling rhizosphere under virgin rotation. Shapes are coloured based on field crop rotation strategy: continuous cropping (orange) and virgin rotation (blue). Shapes indicate compartment: bulk soil (circles), rhizosphere soil (triangles), and roots (squares). Error bars denote a 95% confidence interval around the mean.

**Supplementary Figure S7.**
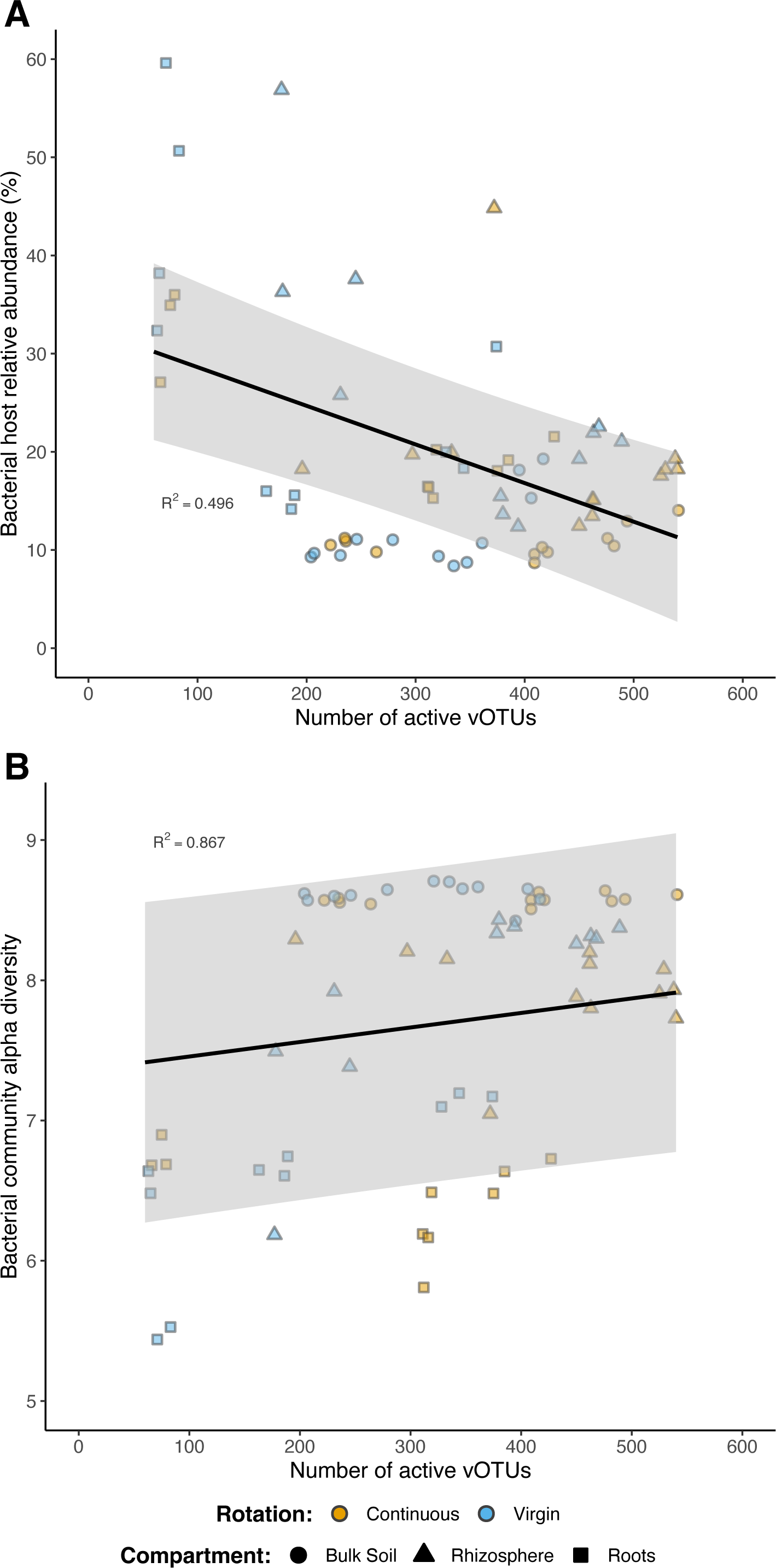
Linear relationships between the number of active vOTUs detected and. **A** Summed host abundance, and **B** Bacterial community alpha diversity. Linear mixed effect models were run using compartment as a fixed effect. Line indicates model prediction, with grey cloud representing a 95% confidence interval around the predicted values. Shapes are coloured based on field crop rotation strategy: continuous cropping (orange) and virgin rotation (blue). Shapes indicate compartment: bulk soil (circles), rhizosphere soil (triangles), and roots (squares).

**Supplementary Figure S8:**
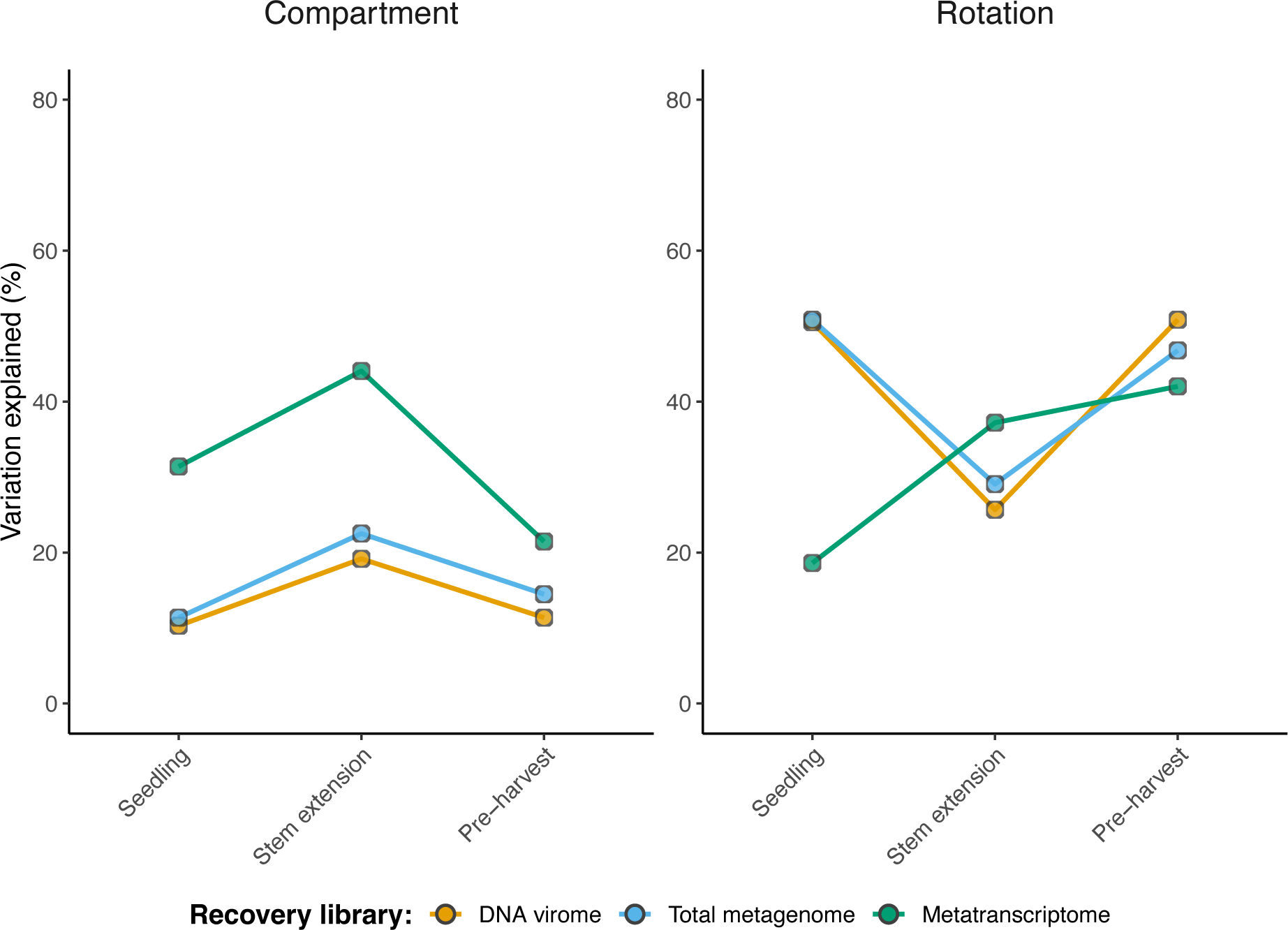
Variation in dsDNA vOTU activity explained by crop rotation and soil compartment by recovery library. PERMANOVA results describe the variance explained by crop rotation and soil compartment, respectively, across growth stages. Points are coloured based on viral activity from vOTUs recovered from the DNA virome (orange; seedling, n = 1,741; stem extension, n = 1,465; pre-harvest, n = 3,140), total metagenome (blue; seedling, n = 368; stem extension, n = 1,379; pre-harvest, n = 1,772), and metatranscriptome (green; seedling, n = 3,844; stem extension, n = 5,078; pre-harvest, n = 5,653).

**Supplementary Figure S9:**
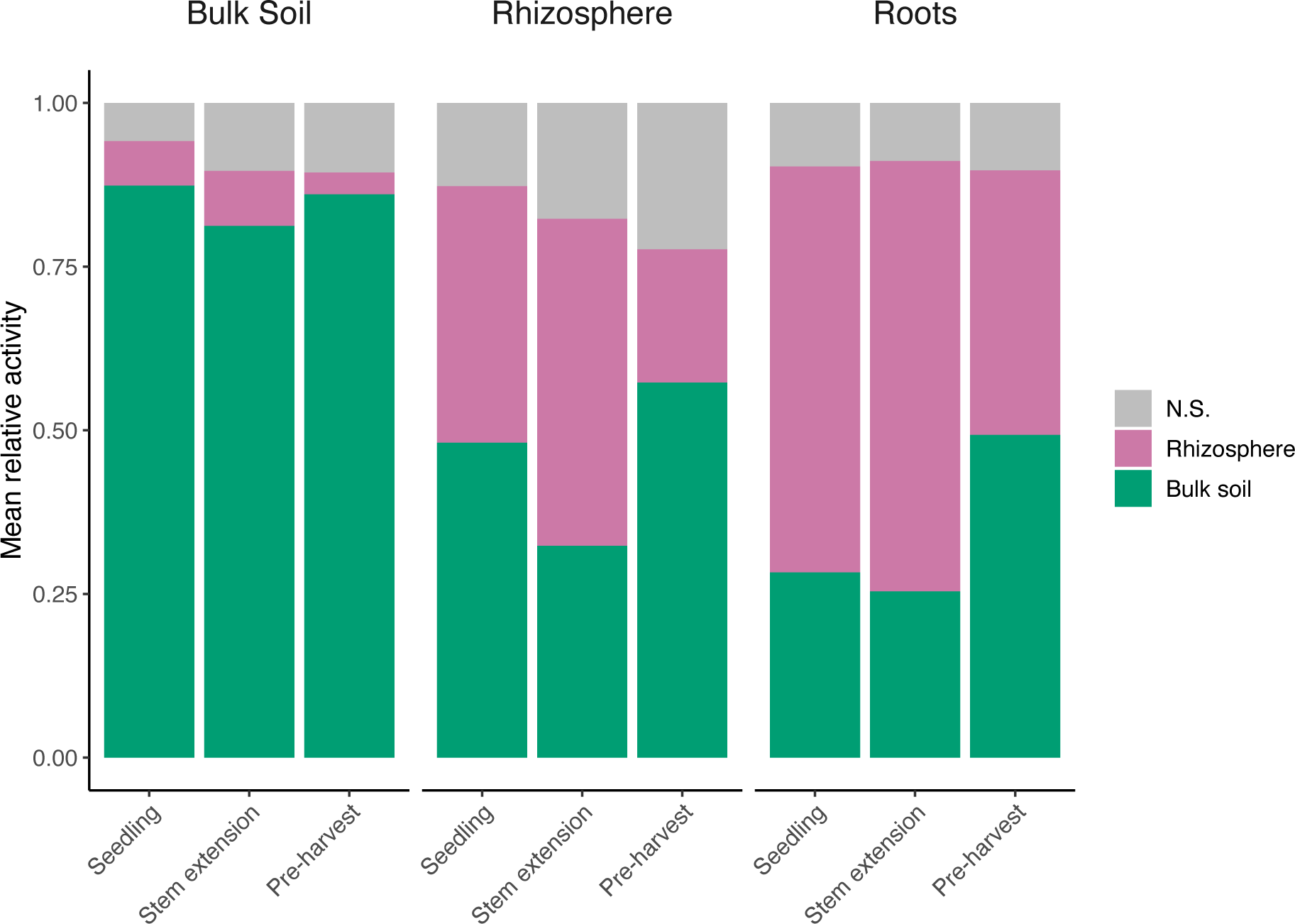
Summed mean relative compartment-enriched viral activity. **A** Relative compartment-enriched viral activity in each root/soil compartment, across growth stages. Colour indicates soil compartment enrichment: N.S. (non-significant; grey), in rhizosphere soil (pink), and in bulk soil (green).

**Supplementary Figure S10:**
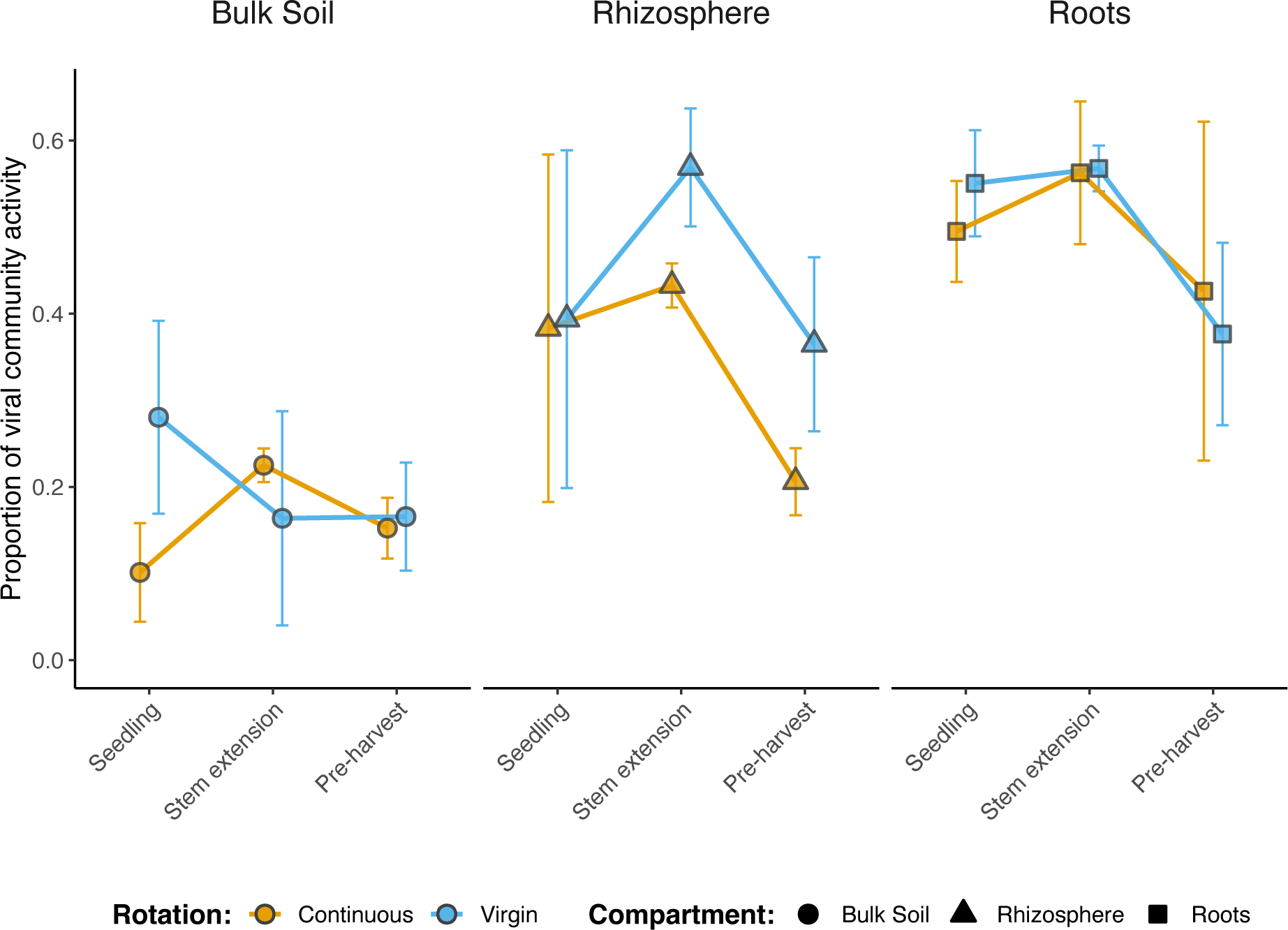
Summed mean relative viral metabolic activity. Proportion of viral community activity represented by viral-encoded metabolic genes in each root/soil compartment, across growth stages. Shapes are coloured based on field crop rotation strategy: continuous cropping (orange) and virgin rotation (blue). Shapes indicate compartment: bulk soil (circles), rhizosphere soil (triangles), and roots (squares). Error bars denote a 95% confidence interval around the mean.

**Supplementary Figure S11:**
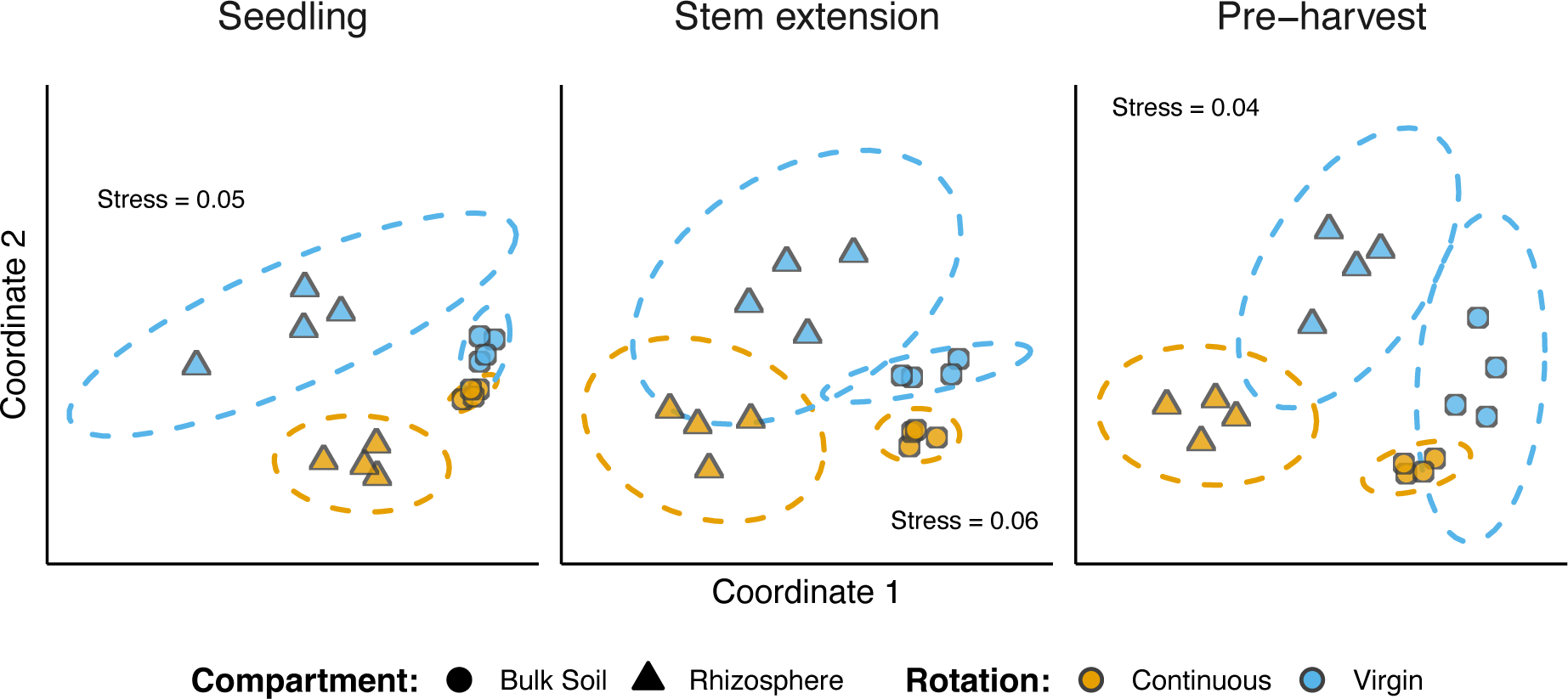
Beta diversity in bacterial community composition. Non-metric multidimensional scaling (NMDS) ordination plots, representing the dissimilarities between community compositions, for each growth stage. Ordinations represent community compositions containing 16S rRNA gene OTUs at seedling (n = 27,335), stem extension (n = 28,235), and pre-harvest (n = 28,958). Shapes are coloured based on field crop rotation strategy: continuous cropping (orange) and virgin rotation (blue). Shapes indicate compartment: bulk soil (triangles) and rhizosphere soil (circles). Stress values associated with two-dimensional ordination are reported for each plot.

## Notes

### Competing Interest Statement

The authors have declared no competing interest.

